# High resolution, serial imaging of early mouse and human liver bud morphogenesis in three dimensions

**DOI:** 10.1101/803478

**Authors:** Ogechi Ogoke, Daniel Guiggey, Tala Mon, Claire Shamul, Shatoni Ross, Saroja Rao, Natesh Parashurama

## Abstract

**Background:** Liver organogenesis has thus far served as a paradigm for solid organ formation. The developing liver bud is a well established model of organogenesis, and murine genetic studies demonstrate key molecules involved in key morphogenetic changes. However, the analysis of the liver bud is typically limited to 2D tissue sections, which precludes extensive visualization, quantitation, and analysis. Further, the lack of human liver bud data has further hindered our understanding of human liver organogenesis. Therefore, new analytical and visualization approaches are needed to elicit further morphogenetic details of liver organogenesis and to elucidate differences between mouse and human liver bud growth.

**Results:** To address this need, we focused on high resolution imaging, visualization, and analysis of early liver growth by using available online databases for both mouse (EMAP, Toronto Phenogenomics center) and human (3D Atlas of Human Embryology), noninvasive multimodality imaging studies of the murine embryo, and mouse/human liver weight data. First, we performed three-dimensional (3D reconstructions) of stacked, digital tissue sections that had been initially segmented for the liver epithelium and the septum transversum mesenchyme (STM). 3D reconstruction of both mouse and human data sets enabled visualization and analysis of the dynamics of liver bud morphogenesis, including hepatic cord formation and remodeling, mechanisms of growth, and liver-epithelial STM interactions. These studies demonstrated potentially under-appreciated mechanisms of growth, including rapid exponential growth that is matched at the earliest stages by STM growth, and unique differences between mouse and human liver bud growth. To gain further insight into the exponential liver bud growth that was observed, we plotted volumetric data from 3D reconstruction together with fetal liver growth data from multimodality (optical projection tomography, magnetic resonance imaging, micro-CT) and liver weight data to compose complete growth curves during mouse (E8.5-E18) and human (day 25-300) liver development. For further analysis, we performed curve fitting and parameter estimation, using Gompertzian models, which enables the comparison between mouse and human liver bud growth, as well as comparisons to processes like liver regeneration. To demonstrate the importance of mesenchyme in rapid liver bud growth and morphogenesis in the human liver bud, we performed functional analysis in which human pluripotent stem cell (hPSC)-derived hepatic organoids were used to model collective migration that occurs in vivo, demonstrating that migration is strongly dependent upon mesenchyme.

**Discussion:** Our data demonstrates improved visualization with 3D images, under-appreciated and potentially new mechanisms of growth, complete liver growth curves with quantitative analysis through embryonic and fetal stages, and a new functional human stem cell-derived liver organoid assay demonstrating mesenchyme-driven collective migration. These data enhance our understanding of liver organogenesis.

## BACKGROUND

The global epidemic of chronic end-stage liver disease has garnered increased interest in liver tissue engineering and liver regenerative medicine, including tools for hepatocyte culture and expansion, pluripotent and adult stem cell biology, tissue chips to replace liver functions, *in vivo* disease modeling, and drug development (Ogoke, Oluwole et al. 2017). The expansion of liver research into these numerous directions has been sustained by fundamental and seminal studies of hepatic, biliary, and hepatic vascular development, reviewed elsewhere (Zorn 2008). These studies detail key molecular and morphogenetic steps in liver organogenesis, which are intricately linked to the liver bud undergoing rapid liver three-dimensional (3D) growth, leading to rapid growth and eventually the largest internal organ in the body. If these 3D morphogenetic steps can be further revealed through imaging and quantitative analysis, this can augment approaches in which human stem cell-derived organoids are employed to model liver organogenesis (Koike, Iwasawa et al. 2019).

Seminal genetic studies have established major stages of early liver bud morphogenesis and the dynamic molecular underpinnings that drive these steps. The liver bud is a transient, multicellular structure that arises during E8.5 in mice and ∼day 25 in humans, on the scale of 100-200 µm in width and length (Si-Tayeb, Lemaigre et al. 2010). At E8.5, the liver bud contains the liver diverticulum, a ventral out-pocketing of the single cell-layered gut tube, surrounded by a layer of endothelial cells, which itself is surrounded ventrally by the septum transversum mesenchyme (STM) (Si-Tayeb, Lemaigre et al. 2010). At E9.0, the pre-hepatic liver epithelium transitions to a pseudostratified epithelium, and thickens in response to cardiac patterning of foregut endoderm via FGF2 (Ledouarin 1963);(Chen, Tillberg et al. 2015) STM patterning of hepatic endoderm via BMP4 (Rossi, Dunn et al. 2001), and endothelial cells patterning of adjacent foregut endoderm towards hepatic endoderm (Matsumoto, Yoshitomi et al. 2001). Genetic studies demonstrate that transcriptional changes correlate with morphogenetic changes. For example, Foxa2, the first known pioneer transcription factor, is upregulated in hepatic endoderm and primes silent liver genes within hepatic endoderm prior to overt morphogenesis (Lee, Friedman et al. 2005). Hex expression occurs during pseudostratification, prior to hepatic cord formation, which is essential for rapid liver growth (Bort, Signore et al. 2006);(Enzan, Himeno et al. 1997). These hepatic cords form at E9.5 (23-25 somite stage) and are defined as finger-like projections that protrude into the STM, express the master transcription factors Hex (Martinez Barbera, Clements et al. 2000), Prox1 (Sosa-Pineda, Wigle et al. 2000), and Tbx3 (Ludtke, Christoffels et al. 2009), and undergo both EMT (Bort, Signore et al. 2006) and collective cell migration (Cascio and Zaret 1991). These studies collectively demonstrate that hepatic cords form rapidly and have a short half-life. Despite their importance, existing histological studies demonstrate hepatic cords with a wide range of geometry, location, and thickness (Matsumoto, Yoshitomi et al. 2001); (Suzuki, Kanai et al. 2006). How these signaling molecules travel in 3D to activate transcription factors and effect 3D morphogenesis is not as well understood. To our knowledge, the hepatic cords have not been imaged/visualized in 3D at high spatial resolution in murine or human liver bud.

The various mesenchymal elements within the liver bud bear a large role in early 3D fetal liver growth and morphogenesis, as borne out by recent scRNA-seq studies (Lotto, Drissler et al. 2020). The STM envelopes the early liver diverticulum in 3D and mediates growth by soluble factor-based induction, leading to transduction of intracellular signals and transcriptional activation. For example, genetic studies of BMP4, Smad2/3 and neurturin (all part of the TGFß superfamily) (Weinstein, Monga et al. 2001); (Tatsumi, Miki et al. 2007), HEP growth factor (HGF) (Schmidt, Bladt et al. 1995), and ephrins (Cayuso, Dzementsei et al. 2016), all demonstrate a role for the inductive properties of STM. Smad 2/3 (TGFß superfamily) haplo-deficient embryos exhibit severely disrupted liver growth by E14.5, due to both loss of ß1-integrin (a known TGFß, target) expression, and mislocalized E-cadherin expression (Weinstein, Monga et al. 2001). HGF knockout studies are mediated by a lack of c-met receptor engagement in hepatoblasts (Schmidt, Bladt et al. 1995). Further, GATA4 -/- embryos lack STM, hepatic cord formation, and subsequent liver development (Watt, Zhao et al. 2007). Knockout of Hlx1, which is strongly expressed within the STM, does not affect early liver specification and hepatic cord formation, but greatly affects 3D liver expansion (Lints, Hartley et al. 1996). Finally, ARF6 knockout disrupts cell membrane polarization, and receptor recycling in liver epithelia (Suzuki, Kanai et al. 2006). In addition to the STM, hematopoietic stem cells, which seed the fetal liver at ∼E11.0, and at 5-6 weeks in human liver, play a role in liver growth. The absence of fetal liver hematopoiesis leads to complex liver morphological effects with mixed phenotypes (Yagi, Deguchi et al. 1998), while affecting liver maturation via the cytokine Oncostatin M (OSM) (Yoshimura, Ichihara et al. 1996), (Kamiya, Kinoshita et al. 1999). Finally, endothelial cells within the developing sinusoids also play a key role in inductive liver growth (Collardeau-Frachon and Scoazec 2008). The absence of endothelial cells in flk -/- mice inhibits liver growth, and *in vivo* studies of explanted mouse liver bud demonstrate that endothelial-lined, branched microvasculature induces 3D liver bud outgrowth even in the absence of blood flow (Matsumoto, Yoshitomi et al. 2001). Moreover, VEGF-A, HGF, IL-6, and Wnt are collectively secreted from sinusoidal endothelial cells and signal to HEPs *in vivo* (LeCouter, Moritz et al. 2003). Taken together, these studies convey the strong inductive, role mesenchyme in 3D fetal liver growth and organogenesis, although imaging/visualization mesenchyme during early liver bud growth has not been achieved (Gordillo, Evans et al. 2015);(Tanimizu and Mitaka 2017);(Zorn 2008);(Ober and Lemaigre 2018). How the STM integrates with the liver tissue in 3D and enables localized collective migration is less well known.

Despite the significance of 3D liver morphogenesis, there have been a lack of studies that determine morphogenetic features in 3D. Established techniques include immunohistochemistry, but this only enables 2D rather than 3D visualization of structure, is highly dependent upon tissue preparation and sectioning angle, and exhibits inter-subject variability (Matsumoto, Yoshitomi et al. 2001, Suzuki, Kanai et al. 2006). To assess human fetal liver growth, scientists have used liver weights or liver morphometry (Szpinda, Paruszewska-Achtel et al. 2015) which don’t enable visualization at the earliest stages, or noninvasive imaging approaches, like ultrasound (Pardi and Cetin 2006, Chen, Tillberg et al. 2015), which lack spatial resolution to image morphogenesis. Noninvasive imaging of liver bud morphogenesis is challenging to perform *in utero* due to tissues, fluids, subject motion, and breathing artifacts. Nonetheless, noninvasive embryo imaging approaches (Nieman, Wong et al. 2011) like light sheet microscopy, (Udan, Piazza et al. 2014), ultrasound (Phoon 2006), optical coherence tomography (OCT) (Syed, Larin et al. 2011), micro-computed tomography (CT) (Wong, van Eede et al. 2015), magnetic resonance imaging (MRI) (Turnbull and Mori 2007), photoacoustic tomography (Turnbull and Mori 2007), optical projection tomography (Lickert, Takeuchi et al. 2004) have been employed. Overall, dynamic imaging has been challenging, and there is still a need for high spatial resolution (∼1-10 µm), dynamic (high temporal resolution), and small field of view imaging to image the earliest dynamic events in liver bud morphogenesis.

The E-mouse atlas project (EMAP) started with a concerted effort for 3D digital visualization of anatomy, histology, and gene expression during embryogenesis (Baldock, Bard et al. 1992), and the 3D Atlas of Human Embryology project followed (de Bakker, de Jong et al. 2016). We hypothesize that these digital resources, as well as others (Toronto Phenogenomics center, and mouse and human liver volume data) could be used to better image and visualize early liver morphogenesis and/or estimate liver volumes. Analysis of the liver bud is limited typically to 2D tissue sections, which limits visualization and quantitation. For this reason, differences between mouse and human liver organogenesis are poorly understood. Therefore, new approaches are needed to elicit further morphogenetic details of liver organogenesis and to compare mouse and human liver bud growth. We focused on high resolution imaging, visualization, and analysis of early liver growth by using available online databases for both mouse (EMAP, Toronto Phenogenomics center) and human (3D Atlas of Human Embryology), noninvasive multimodality imaging studies of the murine embryo, and mouse/human liver weight data. We demonstrate improved high resolution 3D imaging, under-appreciated and potentially new mechanisms of growth, and complete liver growth curves with quantitative analysis. We believe that this will improve our understanding of liver organogenesis. This 3D visualization enables further investigation into the multicellular and mesenchyme specific signals necessary for normal liver organogenesis and microarchitecture (Watt, Zhao et al. 2007);(Zhao, Watt et al. 2005);(Margagliotti, Clotman et al. 2007).

## METHODS

### Mouse liver development data visualization, three-dimensional reconstruction, and analysis

Aligned and registered image slices from each desired time point were downloaded from eMouseAtlas database and e-Mouse Atlas Project (EMAP, emouseatlas.org), (Baldock, Bard et al. 1992) in WLZ format (Richardson, Venkataraman et al. 2014). An outline of the process of data acquisition and analysis is shown (**Figure 1**). In this database, embryos were prepared by fixation, clearing, and embedded in wax, sectioned, stained and mounted on slides. Specimens ranged from 244 to 869 slices with 4 µm per pixel in the x and y dimensions, and 7 µm in the z dimension. The WLZ files were developed from H&E stains of mouse embryos. Each time point was based on one specimen, for a total of (n = 11) mice. Slices were converted to NII format which preserves the scaling of micrometers to pixels for import into ImageJ (https://imagej.nih.gov/). We first located the gut tube within each mouse as a hollow void, extending throughout the specimen. The liver bud starts as the liver diverticulum which is the ventral outpocketing of the gut tube found by locating the liver diverticulum (denoted in the software) surrounded by mesenchyme. We confirmed this is the liver bud by distinguishing it from the lung bud (superior) and ventral and or dorsal pancreatic buds (inferior). A distinct border can be seen of thickened tissue that was identified to be the walls of the liver diverticulum and it and everything inside leading up next to the gut tube was segmented as the liver. The STM is considered to be the surrounding tissue around the thickened epithelial evident from the H&E slides.

**Figure 1.**
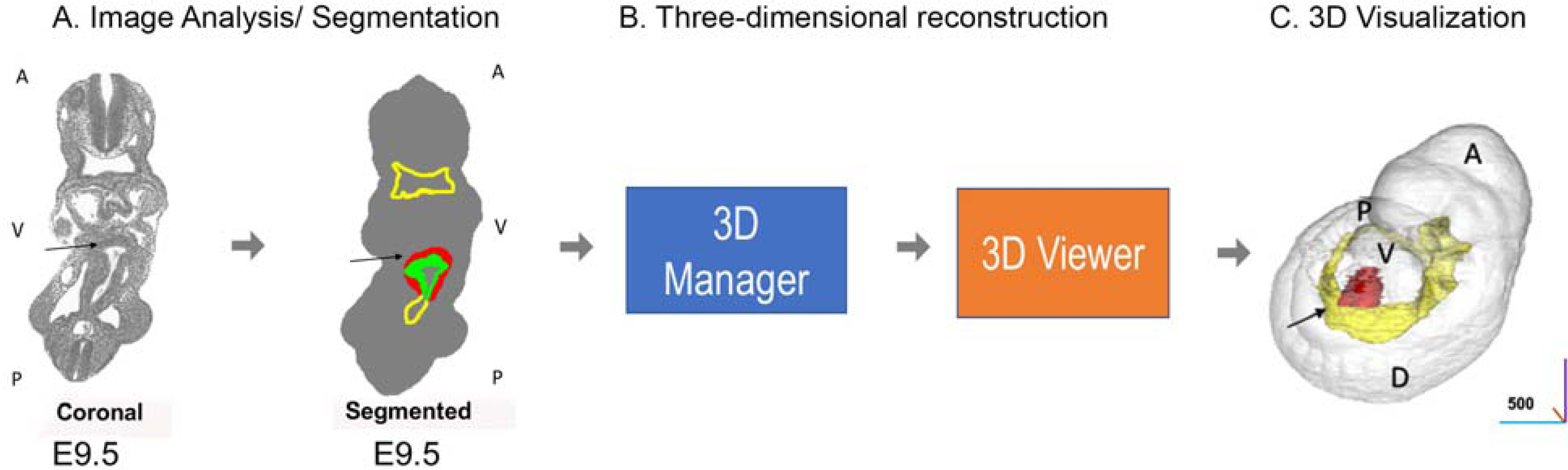
Figure 1. Data processing for 3D reconstruction of mouse liver bud. Listed are the steps in acquiring imaging data, segmenting data, and performing 3D reconstruction. 1) Access emouseatlas.org, mouse anatomy atlas, 2) Choose dataset and time point, 3) Click on download, obtain zip file wlz format of the embryo, 4) Download Woolz from EMAP/Analysis Tools & Resources/Software in the main menu bar 5) Use WlzExtFFConvert from the command line to convert wlz to nii format 6) Import nii format into ImageJ. A similar process was used for human liver as described in methods. A) At this stage, a cleared, sectioned H & E stained embryo is visualized.The image is opened in “Segmentation Editor” and the key parts of the liver bud, mesenchyme, and gut tube are segmented using the mouse atlas with unique colors, using previous liver bud papers or mouse altals as a guide. The label file is saved. B) The 3D Manger (plugin) is used to visualize 3D structure (3D Viewer) and 3D measure is used to perform analysis (i.e. volume). C) 3D reconstruction is performed using 3D viewer with a smoothness factor of 10. Data was checked against mouse atlas and other studies of liver development to ensure accuracy. Scale bar unit is in microns.

We referenced published immunofluorescence of liver (stained for Hnf4a, and Hex) bud tissue to determine relative tissues boundaries of mesenchyme and developing liver bud (Margagliotti, Clotman et al. 2008);(Bort, Signore et al. 2006), existing textbooks and online resources of mouse and human developmental biology to color-code our data sets and reflect accurately *in vivo* representation, (**Figure 1-2, Supplemental Figures 1-3)** Therefore, using Segmentation Editor, pre-tagged developing liver, septum transverse mesenchyme (STM), and the gut tube, were segmented arbitrarily into green, red, and yellow, respectively. E8.5 contained only pre-segmented areas of the liver diverticulum and gut tube, but not the STM. Each 3D object was generated from different EmouseAtlas specimens (EMA). For the E8.5 mouse Theiler Stage (TS) 13, EMA13 was used, at E9.0 (TS14), EMA14 was used, at E9.5 (TS15), EMA28 was used, and at E10.0 (TS16), EMA38 was used. In these specimens, 3D reconstruction of the existing liver bud was performed using the 3D data set which enables the selection of gut tube endoderm, hepatic endoderm, and STM. 3D imaging of the 3D reconstructed mouse liver bud within the whole embryo was unfortunately unable to illustrate significant images of liver bud morphogenesis (**Figure 1-2, Supplemental Figure 1-3**). To focus the field of view to the liver bud only, the embryo was removed from view. When the STM or gut tube was obscuring the liver bud, the opacity of these tissues was reduced to 50%. Light and shading were at default settings and movies were recorded using the built-in tool of ImageJ 3D Viewer. Movies were created by 2 frames per degree of rotation and exported at 7 fps at a resolution of 512 x 512 pixels.

**Figure 2.**
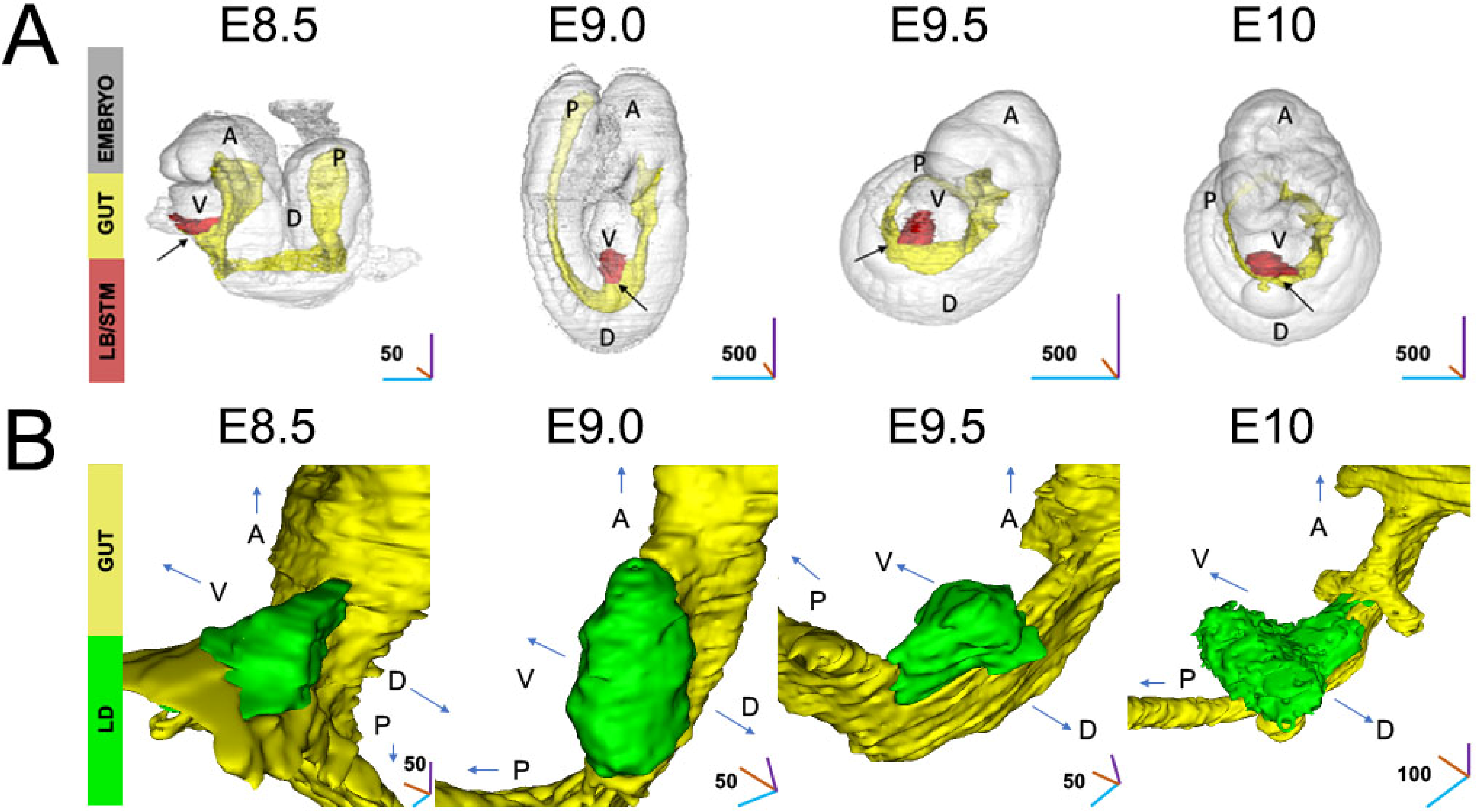
3D reconstruction and visualization of the developing murine liver bud. To dynamically visualize morphogenetic stages during mouse liver bud formation, 3D datasets, consisting of cleared hematoxylin and eosin (H& E) tissue sections from mouse development were used. Datasets were reformatted, imported, segmented, and reconstructed in 3D. Ventral views are shown. A = anterior, V = ventral, P = posterior. Arrows depict directions, gut tube (yellow), the liver bud (green). One specimen was used for each time point. A) Whole body 3D reconstruction of the developing embryo at transient times of development (E8.5, E9.0, E9.5,E10). The major components of the murine liver bud are highlighted in separate colors. Gut tube (yellow), and Liver bud (LB) and septum transversum mesenchyme (STM) in red. Arrows (black) indicate location of the LB/STM. Scale bar in first panel is in µm. Scale bar in next three panels is in microns. B) 3D reconstruction of the posterior foregut focused on the liver murine liver bud, at E8.5 from Theiler Stage (TS) 13. Liver epithelium initially appears as a triangular sheet of cells with a base of 25 µm At E9.0 (TS14)gut tube has closed with the liver bud, forming an elliptical shape tube-like structure with long axis in the cranial-anterior direction. For E9.5 (TS15) the liver bud remodels its shape, with an elliptical long axis in the lateral direction. At E10.0 (TS16). Liver bud remodeling with an elliptical long axis extending bi-laterally. At this stage, the axis has completely changed from E9.0. The liver surface appears more roughened.

For calculations of mouse liver volume, we also downloaded volumetric data for MRI and optical projection tomography (OPT) from the eMouseAtlas database and e-Mouse Atlas Project (EMAP, emouseatlas.org) (Baldock, Bard et al. 1992) in WLZ format (Richardson, Venkataraman et al. 2014). This OPT volume data included E9.5 (n = 2), E10 (n = 1), E10.5 (n = 2), E11.0 (n = 1), E11.5 (n = 1) and the MRI volume data included E14.0 (n = 1) and E18.0 (n = 1). In addition to the mouse embryo image stacks obtained from the e-Mouse atlas, we also acquired volumetric data from the Toronto Centre for Chemogenomic Mouse Imaging Centre for E11.5, E12, E13, and E14 (http://www.mouseimaging.ca/). This data was obtained using optical projection tomography (OPT) and computed tomography (CT). For OPT data, volume data was obtained for E11.5 (n = 1), E12 (n = 1), E13 (n = 1), and E14 (n = 1). For CT data, volume data was obtained for E15.5 (n = 35). The mouse number obtained from each analysis approach (**Table I**) and the voxel dimensions (**Table II**) are shown. The mouse strains were C57/BL6. Using the same procedure, we performed 3D reconstruction followed by calculation of the volume as above.

**Table I:**
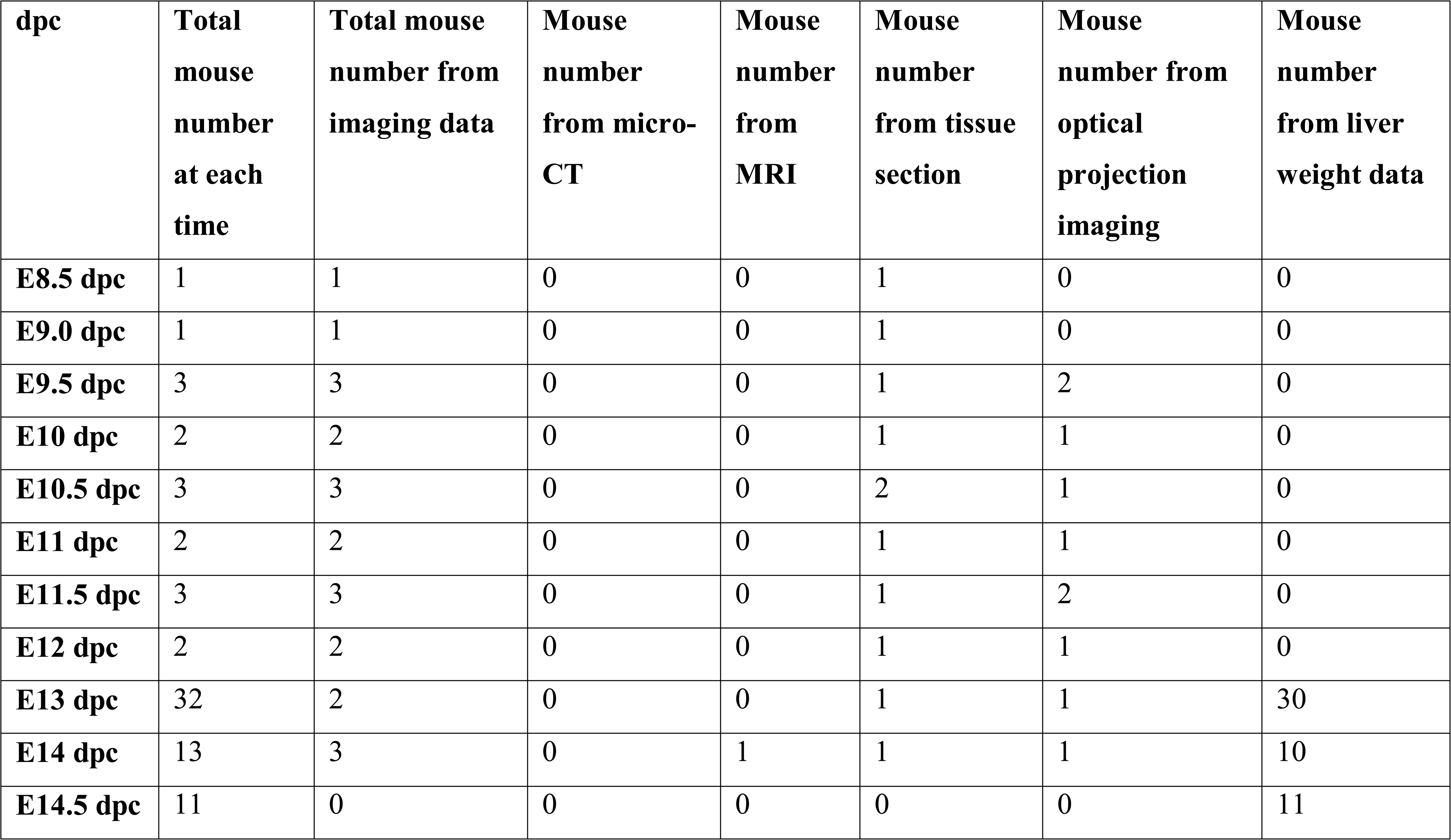
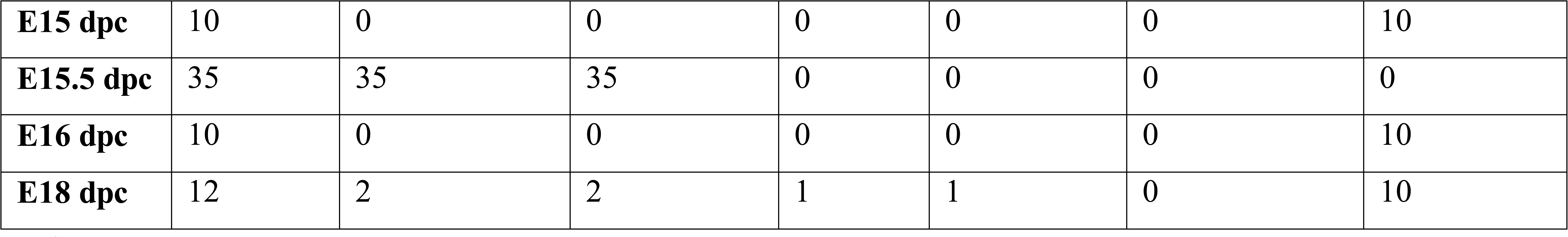
Total mouse number obtained from imaging modalities and liver weights.

**Table II.**
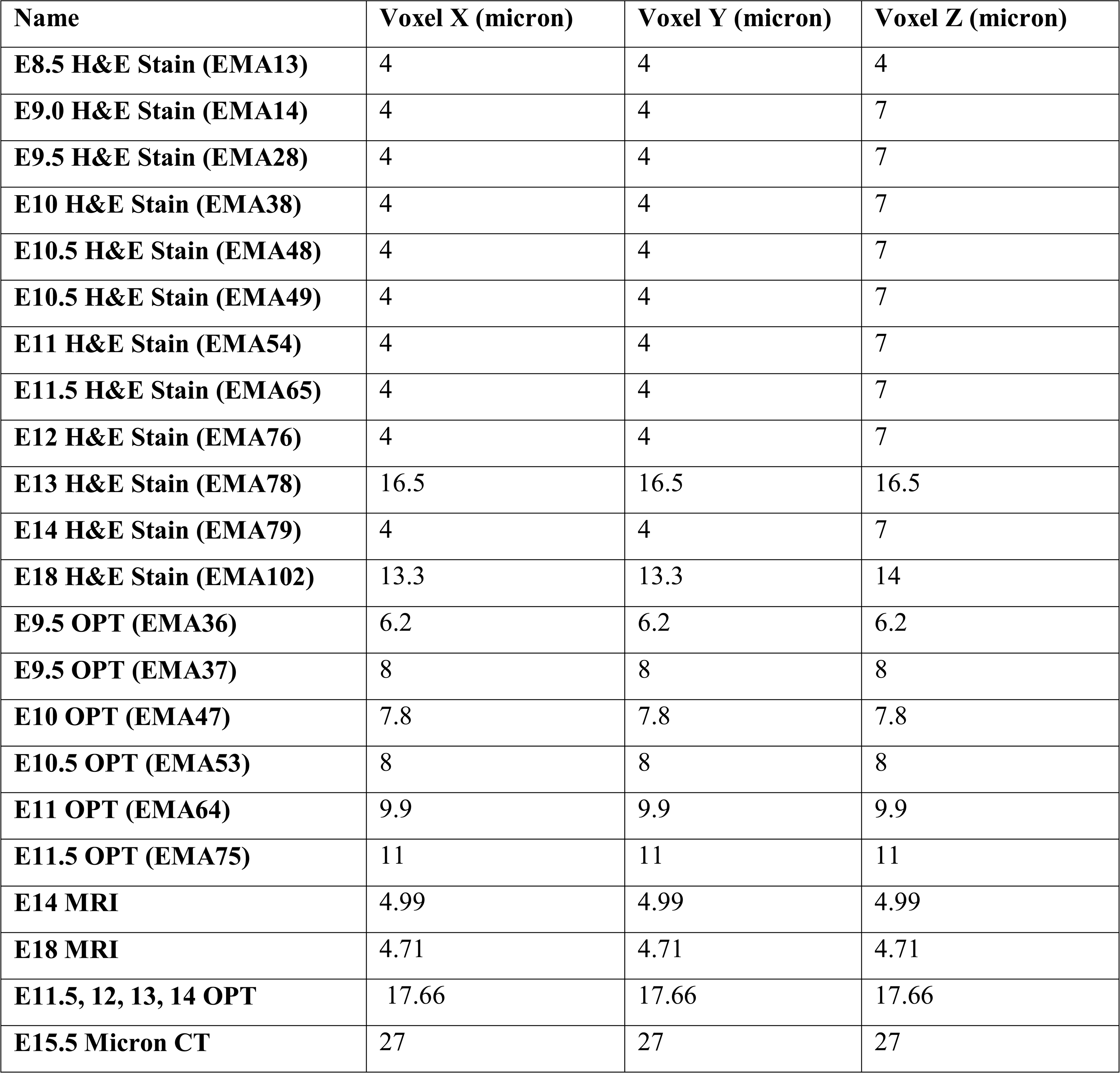
Voxel size information for each mouse data set.

### 3D reconstruction and visualization of human liver bud

For the human development, gray image slices of the aligned and registered H&E stains were downloaded from the 3D Atlas of Human Embryology (3dembryoatlas.com) along with the label files (de Bakker, de Jong et al. 2012). 3D Atlas of Human Embryology utilizes about 15,000 sections, which were digitized and aligned with the software package Amira to create image stacks. Gray and label files are imported into ImageJ using the TrakEM2 plugin and the scaling of micrometers to pixels is set using values provided by the authors. The STM was segmented manually, and visualized from histological sections, whereas for the developing liver and the gut tube we used the pre-existing segmentation. All segmented regions are exported as image stacks in TIFF format. 3D reconstruction was performed by importing the stacks into the 3D Viewer plugin with a smoothness factor of 10. Pixel values for each time point (Carnegie stage) are listed (**Table III**). Movies were created by 2 frames per degree of rotation and exported at 7 fps at a resolution of 512 x 512 pixels.

**Table III.**
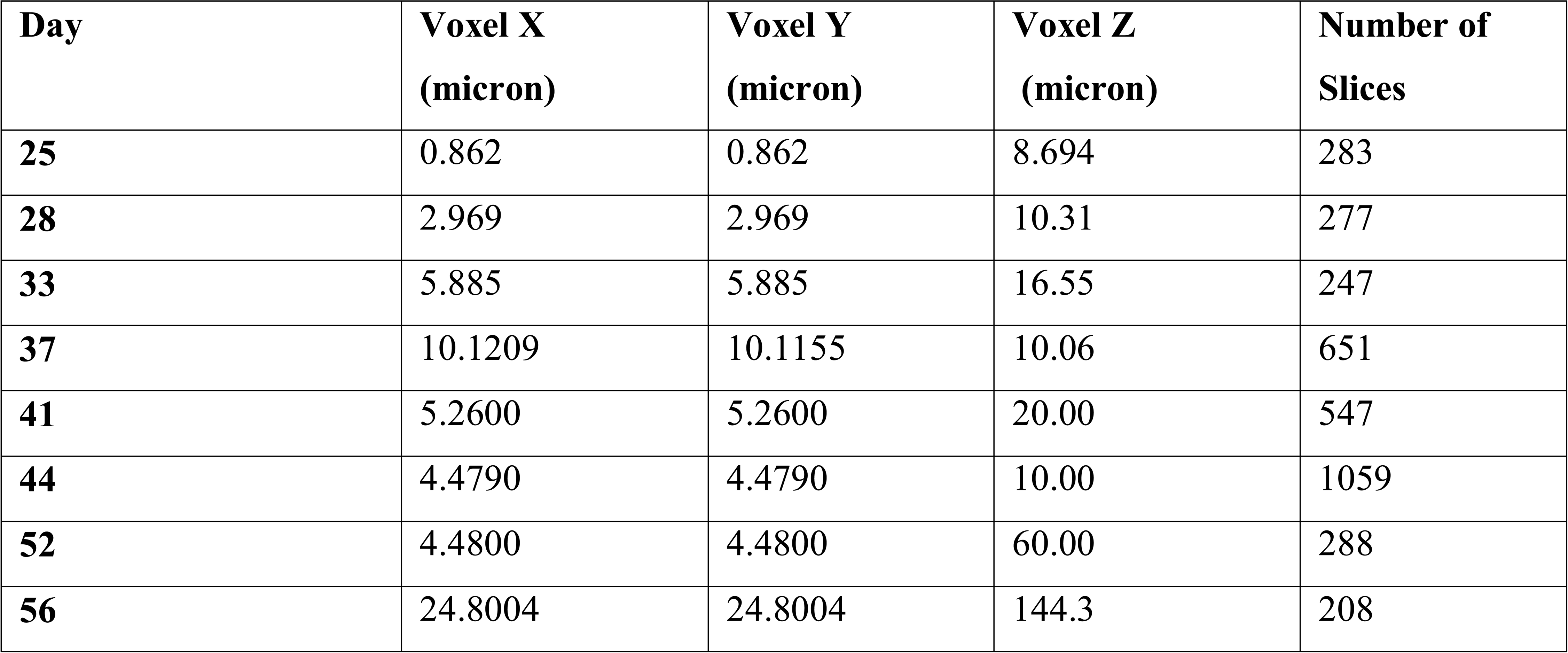
Voxel size for human studies.

### Calculation of mouse and human volumes from liver weights

We also obtained mouse and human liver weight data and estimated volumes from weights. Mouse liver weight data was acquired for E13, E14, E15, and E18 (Nomura 1976). Additional mouse liver weight data was acquired from (Kamei, Tsutsumi et al. 1999) and (Song, Luo et al. 2008)for E13 and E14.5 respectively.

Human fetal volumes (n=69) were obtained (Szpinda, Paruszewska-Achtel et al. 2015) in addition to liver weight data (n=861) (Man, Hutchinson et al. 2016). The human liver weight data were obtained from human fetuses from (Man, Hutchinson et al. 2016) for days 84 through 294. To account for the growth-restricted liver weights in (Man, Hutchinson et al. 2016), the bottom 10^th^ percentile of the data was removed. The bottom 10^th^ percentile was determined by finding the 4^th^ order polynomial regression line for the liver weight data to find points furthest from this curve. Since the noise increases proportionally with the volume, the error for each data point was adjusted by the factor 1/*V*. The 96 points out of the 957 data points with the highest negative error were removed from further analysis. By assuming the liver has a density of water (∼1 gm/cm^3^), the liver volume was able to be estimated for each liver weight.

### Calculation of volumes and surface area of embryonic structures

Using the 3D Manager plugin within ImageJ, volume and surface area were measured with the Measure 3D option after loading the image stacks.

### Estimation of cellular density in the gut tube, liver, and STM

For each region, the original histology was examined and imported into ImageJ. Random sections of each region were selected, and the number of cells was counted. This count was divided by the area of the sections. It was assumed that the height of each cell was 10 µm and further divided by a height estimation of 10 µm, to give a value of cells per cubic micron, a single cell volume of 1000 um^3^, as done previously (Semeraro, Cardinale et al. 2013).

### Statistical methods, linear regression analysis and Gompertz modeling

Linear, exponential, logistic, and Gompertz regression models were tested as models for our liver development growth data. The Gompertz model was found to produce the highest coefficient of determination (r^2^) and lowest root mean square error (RMSE) for our data and thus was chosen as the model to fit the data. Gompertz has been used as a model for tumor growth (Yang, Gao et al. 2020);(Mehrara, Forssell-Aronsson et al. 2013). The Gompertz model can be described as 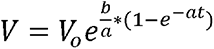 Where *V_o_* is the initial volume, b and a are constants, and V is volume with respect to time t. In this analysis, volume estimates for mouse and human livers were plotted against dpc and examined by non-linear regression analysis against Gompertz growth curve. GraphPad Prism software (GraphPad Software Inc., San Diego, CA) was used to estimate the constants to fit the data as well as find r^2^-values, RMSE, p-values, and confidence intervals. The parameters were estimated by least squares regression. The sum-of-squares of the distances from the curve was also weighted by 1/*V*, since the magnitude of error was found to increase proportionally with the magnitude of the volume. This analysis was done for both mouse and human liver growth. The mouse analysis was done with all the multimodality volume estimates (n=59) as well as with the estimated volumes from liver weight data Nomura et al. (n=10) (Nomura 1976), Kamei et al. (n=20) (Kamei, Tsutsumi et al. 1999), and Song et al. (n=11) (Song, Luo et al. 2008). The human embryo 3D reconstruction volume estimates (n=9) were analyzed together with measured human fetal volumes (n=69) (Szpinda, Paruszewska-Achtel et al. 2015) as well as liver weight data (n=861) (Man, Hutchinson et al. 2016).

### Reagents/materials

DMEM (Thermofisher, Cat. #: 10569010), FBS (ThermoFisher, Cat. #: A3160701), Pen-Strep (10000 U/ml) (Thermofisher, 15140122), 0.05% Trypsin-EDTA (Cat. #: 25300062), Matrigel (MG) Growth factor-free (Cat. #: 40230), 384-well round bottom, ultra-low attachment spheroid microplates (Corning, Cat. #:3830) was purchased from Corning. Aurum Total RNA Mini Kit (Cat. #: 7326820), DNase I (Cat. #: 7326828), iTaq Universal SYBR Green Supermix (Cat. #: 1725121), and iScript cDNA Synthesis Kit (Cat. #: 1708891) were purchased from Bio-Rad. Tissue Culture Treated 24-well plate (LPS Cat. #: 702001), 75-cm^2^ Polystyrene tissue Culture-Treated Flasks (LPS Cat. #:708003), PCR Plate Covers (LPS Cat. #: HOTS-100) were purchased from Laboratory Products Sales (LPS). All PCR primers were purchased from either Integrated DNA technologies (IDT), Sigma Aldrich, or Thermofisher. Human foreskin fibroblasts were a obtained as a kind gift from Dr. Stelios Andreadis’ lab at the University at Buffalo, State University of New York.

### Feeder-free culture (maintenance) of human pluripotent stem cells

We performed experiments using a commercially available induced pluripotent stem cell (iPSC) line (BXS0114 ACS1028 female (ATCC). Human pluripotent stem cells (hPSC) were cultivated at 90% N_2_, 5% O_2_, 5% CO_2_, (Tri-gas HERAcell VIOS 160i CO2 incubators) using room temperature mTESR1 medium on 6-well tissue culture-treated plates, coated with 1:15 diluted (in DMEM) growth factor free-matrigel (MG). Wells were coated with MG by adding 1 mL of diluted MG per well of a 6-well plate, and incubating for 1.5-2 hrs. at 37°C. Excess dilute MG was then removed and the wells were washed with PBS. Cell culture medium was changed every other day. For passaging, the mTESR1 serum-free maintenance medium was removed, 1 mL of gentle cell dissociation reagent (GDR) (Stem Cell Technologies) was added for 10-15 minutes, and single cells or clumps of cells were harvested from the dish. Cells were centrifuged 3 minutes at 800-1000 RPM (Eppendorf 5810 table top centrifuge) and resuspended. Cells were frozen in mTESR1 medium + 5% DMSO at −80°C overnight, followed by liquid nitrogen cryostorage. Passage number varied between 15-35 for all experiments.

### Preparation of stem cell differentiation medium (SFD medium)

Prepared as stated in another publication (Warren et al. 2020). Basal differentiation media for hPSC differentiation to endoderm, gut tube, and liver is comprised of SFD, a serum-free, defined medium based upon additional mouse and human stem cell studies (Gadue, Huber et al. 2006). SFD medium is composed of RPMI basal media containing (75 % IMDM, 25% Ham’s F12, 0.5% N2 Supplement, 0.5% B27 supplement, 2.5 ng/ml FGF2, 1% Penicillin + Streptomycin, 0.05% Bovine Serum Albumin, 2 mM Glutamine, 0.5mM Ascorbic Acid, and 0.4mM Monothioglycerol (MTG)).

### Generation of hPSC-derived definitive endoderm

An endoderm protocol was developed specifically for improved survival at low oxygen and improved morphology. In RPMI medium with 1x B27 (no insulin) and 0.2% KO serum, definitive endoderm from iPSC cells was induced with Activin A (100 ng/ml) and CHIR (3 µM) for 1 day, followed by Activin A (100 ng/ml) for 3 more days. Medium was changed daily and 500-750 µL medium was used per well. The 0.2% KO serum was added for improved viability at 5% O_2_, and higher seeding densities improved culture morphology.

### Generation of hPSC-derived hepatic progenitor cells

To promote hepatic differentiation with growth factors, we adopted a protocol from the literature (Takebe, Sekine et al. 2013). Briefly, on day 4 of definitive endoderm induction, Bone morphogenetic protein 4 (BMP4, 10 ng/ml), and fibroblast growth factor 2 (FGF2, 20 ng/ml) were added from days 4-9, and Hepatocyte growth factor (HGF, 10 ng/ml), Oncostatin M (20 ng/ml), and Dexamethasone (100 uM) were added from days 9-12. Cells were assayed by qRT-PCR on day 12.

### 384-well plate organoid formation

Hepatic progenitor cells, were obtained from hPSCs differentiated on feeder free Matrigel coated plates under low oxygen. To make endoderm spheroids, differentiation media was aspirated from each well of the plate and Accutase was added to each well and allowed to incubate on the cells for 10 minutes at room temperature. Following this, cells suspended in Accutase media were collected into a 15 mL tube and spun down at 1000 rpm for 5 min to pellet the cells. The Accutase was then removed and the cells were re-suspended in serum free differentiation media (SFD) containing Rho-kinas inhibitor (1:1000) and knockout serum (KOSR, 20%) and counted using a hemocytometer. The total number of cells needed per well was 20,000 so adjustments were done to have an appropriate number of cells for seeding. Cells compacted in 24 hours in 384 ultra low attachment, (ULA) round bottom plates. In the case of co-culturing with stromal cells i.e., human foreskin fibroblasts (HFF), these fibroblasts were first harvested from T-75 cells and counted and adjusted in concentration to 1×10^6^/mL in supplemented DMEM. These cells were then centrifuged and resuspended in ice-cold Matrigel at the same concentration (1×10^6^/mL of Matrigel) before being seeded into the same 384 well plates containing the hepatic organoids in suspension (20 uL, 2 x 10^5^ cells per well).

### Analysis of organoid migration and shape

ImageJ was used to analyze the relative properties of the organoids in culture. Collected images were uploaded to the Image J tool. A global scale bar was used per image uploaded before analysis. The length application in Image J was used to calculate protrusion length of the separate experimental conditions. In addition, the circularity plugin was used to determine the shape alterations of the central organoid over time in culture. Briefly, each organoid image was converted to 8 bit gray scale and then its outline traced to determine the corresponding circularity.

### Phase and brightfield microscopy

For phase microscopy, iPSC derived 2D cells and 3D organoid cultures were imaged with a benchtop microscope specifically a Zeiss Axio fluorescence microscope (SE64, 1344×1024 pixel density) equipped with Axiovision Software (v4) and analyzed using Image J.

### RNA isolation, reverse transcription (RT) and quantitative polymerase chain reaction

Total RNA was purified with Aurum Total Mini Kit (Bio-Rad) using the spin column method with DNase 1 (Bio-Rad, Hercules, CA) (reconstituted in 10mM tris) treatment. RNA concentrations were determined by Nanodrop. RNA was converted to cDNA with an RT reaction using the iScript cDNA Synthesis Kit (Bio-Rad), and the mass of RNA was calculated such that 5 ng RNA per well to be run in the qPCR reaction. The RT reaction was performed using 5 minutes at 25° C, 20 minutes at 46° C, and 1 minute at 95°C. Reactions were then held either at 4° C or on ice. We performed 10 µL qPCR (3 µL primers at a concentration of 0.3 µM, 1 µL nuclease free water, 1 µL cDNA, and 5 µL supermix) reactions with iTaq Universal SYBR Green Supermix (BioRad) in a 96-well PCR plate (LPS). The qPCR reaction was done in a CFX96 Touch Real-Time PCR Detection System (BioRad). The qPCR reaction consisted of polymerase activation and DNA denaturation at 98°C for 30 seconds followed by, 40 to 45 cycles of 98°C for 15 seconds for denaturation and 60° C for 60 seconds for annealing and extension. Melt curve analysis was performed at 65-95° C in 0.5°C increments at 5 seconds/step. Relative, normalized, gene expression was analyzed using the delta-delta-Ct method (Livak and Schmittgen 2001), with three duplicates per gene tested

### Statistics

Data collected was reviewed and analyzed by Microsoft excel. The Student’s t test was used to determine statistical differences between two or more independent groups (P value set at <0.05).

## RESULTS

### 3D serial imaging of early liver bud morphogenesis

To image the murine liver bud at high resolution in 3D, we performed 3D reconstructions of stacked, digital, cleared, hematoxylin and eosin (H& E) tissue sections from mouse development at key time points (E8.5, E9, E9.5, and E10) and analyzed the 3D images. At each time point, we segmented each tissue section, targeting the liver epithelium, gut tube, and STM. Upon analysis, the liver bud epithelium is distinct and clearly visualized (green), and demonstrates the gut tube (yellow), the STM (red), and the liver bud (green) (**Figure 1A, E9.5**). Using these representative, segmented sections, together with segmented images for E8.5, E9.0, E9.5, and E10 (**Supplementary Figs 1-3**), we first reconstructed the entire embryo in 3D, which enabled visualization of the colored liver bud and gut tube (**Figure 2A).** These images clearly demonstrate growth in the embryo and liver bud, but liver epithelial morphogenesis (green) could not be visualized, whereas the STM (red) and gut tube (yellow) are clearly visualized (**Figure 2A**). To visualize the liver bud dynamics over time, we reduced the field of view to reveal only the 3D liver bud epithelium, STM, and gut tube (**Compare with Figure 2B with 2A**). When viewed as part of the gut tube, the E8.5 liver epithelium appears as a triangular sheet of cells (**Figure 2B, E8.5**) with tube-like and flat sheet-like regions, which we estimate to be approximately 295 µm (base) x 288 µm (height) x 75 µm (thickness). Although there is known to be transitional epithelium with varying cell size in the liver epithelium (Sherwood, Chen et al. 2009) (**Supplemental Figure 4, Supplemental Video 1-2**), we estimate approximately 3186 cells in the E8.5 liver bud, assuming a single cell volume of 1000 um^3^, as done previously (Semeraro, Cardinale et al. 2013). Our morphogenesis analysis of 3D liver bud demonstrates an elliptical shape at E9.0 in the cranial/anterior direction, with a rapid transition to an elliptical shape in the lateral direction at E9.5-E10.0 (**Figure 2B**, Compare E9.0, E9.5, and E10).

### Imaging of migrating hepatic cords and sheet-like growth in the developing mouse liver bud

Having imaged liver bud remodeling in 3D, our goal was to visualize and analyze microscopic aspects like the hepatic cords and morphogenetic features. We segmented out the gut tube from the images and focused only on the liver bud. With 3D reconstruction and multiple rotated views of the E9.0 liver bud, we observed numerous miniature finger-like projections with dimensions of 1-10 µm in length/ diameter (**Figure 3A-C (arrows), Supplementary Video 2**). We also observed sheet-like growth, arranged in a series of ridges spaced between 20-30 µm, at E9.0, from the superior to the inferior portion of the liver bud (**Figure 3A-C, Supplementary Video 2**). Since the original specimens range from 244 to 869 aligned (image registered) slices and the z dimension (height) interslice distance 7 µm in the z dimension, these sheet-like ridges are not a result of imaging artifact or artifacts associated with stacking. They are not a result of interpolation since interpolation was not performed, and they are not due to loss of liver bud tissue during sectioning, since images are based on whole embryo tissue slices. Importantly, the sheet-like ridges do not appear within the gut tube, but solely within the liver tissue, further precluding the above mentioned possibilities, and indicating that these morphogenetic structures were liver bud-specific. At E9.5, we observed slightly larger finger-like projections (5-10 µm) (**Figure 3D-G, Supplementary Video 3**), indicating 3D growth, and sheet-like 3D growth arranged in larger ridges at E9.5 compared to E9.0 (**Figure 3E, Supplementary Video 3**). Our data at E10 represents liver bud tissue in which liver epithelial cells (hepatoblasts) and mesenchymal cells intermingle. Because of these, the STM cells are included within the green labeled liver tissue. The E10 liver bud demonstrates numerous interconnections cells that form a sponge-like, trabeculated tissue throughout the liver bud, but not in gut tube (**Figure 3H-K, Supplementary Video 3-5**). To demonstrate variability and repeatability problems in analysis of the mouse liver bud, we demonstrate n = 4 tissue sections from existing literature for both E9.0 and E9.5, which demonstrate inter-study and intra-study variability (**Figures 3L-M**). While these studies demonstrate variability, they also validate 3D reconstruction data. Overall, we demonstrate 3D hepatic cords, 3D sheet-like growth pattern at E9.0-9.5, 3D visualization of trabeculation by E10, and variation present within existing studies.

**Figure 3.**
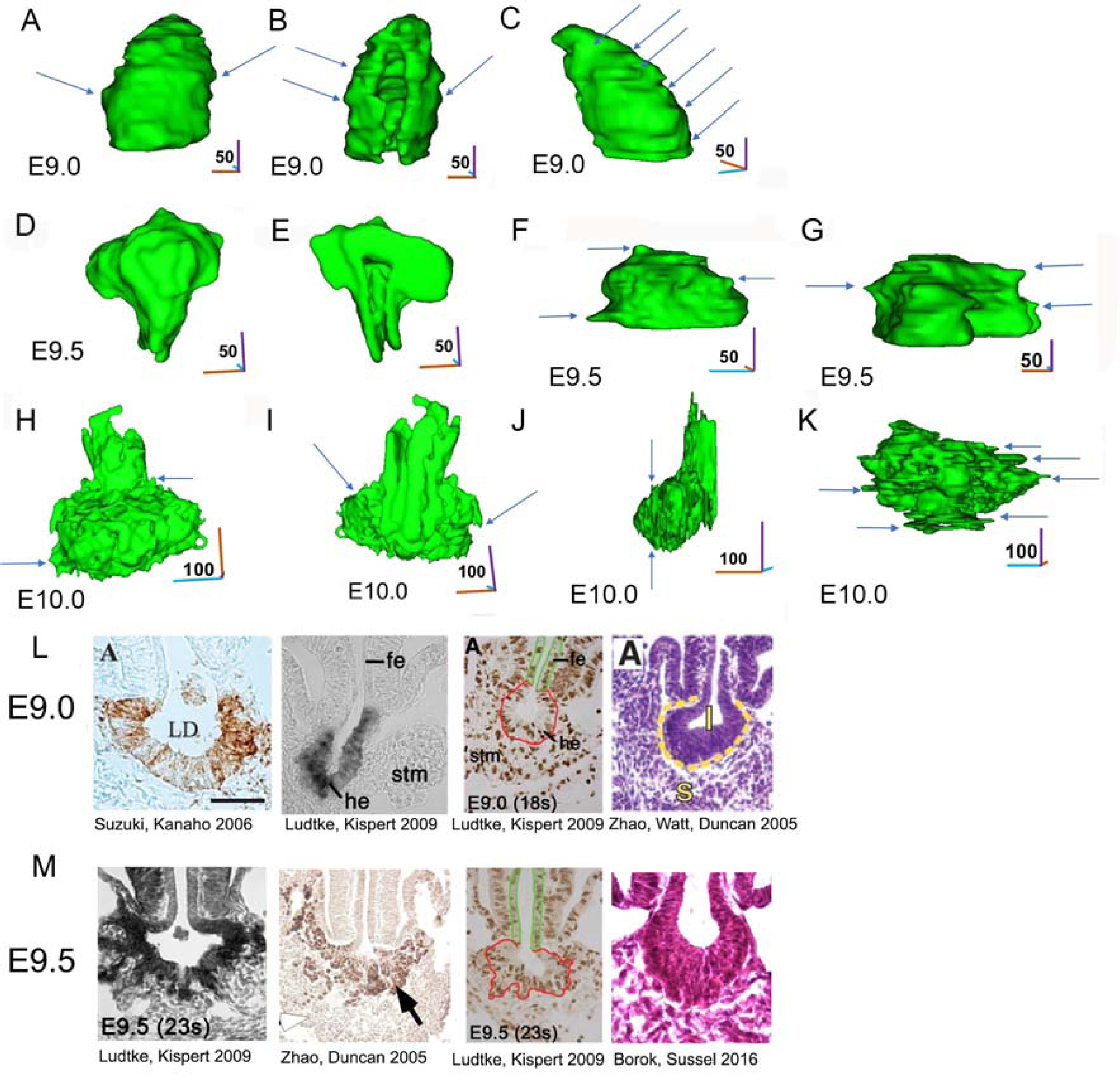
Multiple views of 3D murine liver bud demonstrate dynamic formation of cord-like structures and sheet-like growth. A) 3D reconstruction of the 9.0 murine liver bud, ventral view. Arrows demonstrate multiple early hepatic cords which demonstrate migration in the lateral directions. B) Same structure as A), except dorsal view. The dorsal view demonstrates how the liver bud wraps around the gut tissue (which is absent), with an associated lumen. The associated hepatic cords are depicted by arrows. C) Same structure as A), except right lateral view. In this case, arrows depict not hepatic cords, but sheet-like growth at various levels in the cranial-anterior axis. This sheet-like growth appears most prominently in this view. D) 3D reconstruction of the E9.5 murine liver bud (TS15), ventral view. E) Same structure as D), except dorsal view. F) Same structure as D), except dorsal view, inferior view. Blue arrows depict hepatic cords, while sheet-like growth is not observed. G) Same structure as D), except left lateral view. Blue arrows depict hepatic cords, while orange arrows depict sheet-like growth. H) 3D reconstruction of the E10.0 murine liver bud (TS16). Arrows depict examples of migrating cords. The liver bud now demonstrates a trabeculated tissue with interconnecting open-spaces. I) Same structure as H), except dorsal view J) Same structure as H), except left lateral view, demonstrating trabeculated tissue. Arrows depict potential hepatic cords. K) Same structure as H), except inferior view, demonstrating trabeculated tissue. Arrows depict potential hepatic cords. L) Cross sections of mouse liver bud at E9.0 as reported in the literature. 4 examples given with references below. M) Cross sections of liver bud at E9.5 as reported in the literature. 4 examples given with references below.

### Quantification of 3D liver bud volumetric growth

The liver bud leads to rapid expansion to result in the largest organ in the body, but it has not been formally determined how rapid embryonic liver bud growth (E8.5-11.5) physically matches with fetal liver growth (E12-18). To determine this mathematical and physical relationship, we employed data not only from 3D reconstructed tissue sections, but also from micro CT, OP imaging, MRI, and data from published liver weights (converting liver weight to volume), to validate our approach. To add additional data at multiple time points, we identified studies that had measured normal fetal liver weights (Nomura 1976);(Kamei, Tsutsumi et al. 1999);(Song, Luo et al. 2008). To convert weight data to volume data, we assumed a density equal to water (1 g/mL) (Overmoyer, McLaren et al. 1987); (Xie, Schwen et al. 2016). We performed a growth analysis of the liver by analyzing liver volume growth over time. Combining embryonic and adult data demonstrated a clear, exponential increase in growth over 5 orders of magnitude (**Figure 4A, Table IV**). This data agrees with our estimation of growth using E8.5 to E18 measurements from 2D images (data not shown). To better visualize these growth changes, we normalized the data (**Figure 4B)** and performed a nonlinear regression analysis on both the absolute (R^2^ = 0.85, and P < 0.001, RMSE = 11.42 x 10^9^) and normalized data absolute (R^2^ = 0.85, and P < 0.001, RMSE = 3.29 x 10^4^) with a Gompertz model (**Figures 4C-D**), an approach used widely in tumor growth studies (Yang, Gao et al. 2020); (Mehrara, Forssell-Aronsson et al. 2013). We additionally wanted to estimate cell count to place our results in greater context. We estimated cell number using an estimated cell size of (10^3^ µm^3^) (Semeraro, Cardinale et al. 2013). The cell number was calculated by dividing the liver volumes in our Gompertz model by this estimated cell volume to estimate the number of cells in the liver over time. Interestingly, the cell kinetics model obtained agrees with previous data (Konstantinidis, Giger et al. 2015);(Ema and Nakauchi 2000) for data between E11 and E17 (**Figure 4E**). This indicates that cell proliferation may play a greater role in the increase in cell volume than hypertrophy in liver development. Furthermore histology sections within the fetal liver) this time period between dpc 12 and dpc 16 do not demonstrate noticeable increases in cell size or cellular hypertrophy (**Figure 4F**) (Ayres-Silva Jde, Manso et al. 2011). It is possible that the ability of hepatocytes to regenerate in acute regenerative models exhibits proliferative capacity similar to embryonic liver growth. To this answer question, we compared our Gompertz model to the rapid volumetric growth that during acute liver regeneration (Nevzorova, Tolba et al. 2015). When regeneration data was normalized, and shifted to start at the same time as liver bud growth, growth during liver organogenesis was over 20,000 times the growth during regeneration (**Figure 4G**), suggesting different mechanisms at play. Consistent with this, the volumetric growth rate during acute regeneration was found to be similar to the Gompertzian volumetric model at day 19 to day 23, when liver bud growth rate is decreased by 90% of the initially rapid rate at E8.5 (data not shown). Overall, we demonstrate a model of growth that links embryonic and fetal liver growth which clearly correlates well with existing data. This model was used to estimate cell kinetics over time as well as liver bud growth compared to liver regeneration.

**Figure 4.**
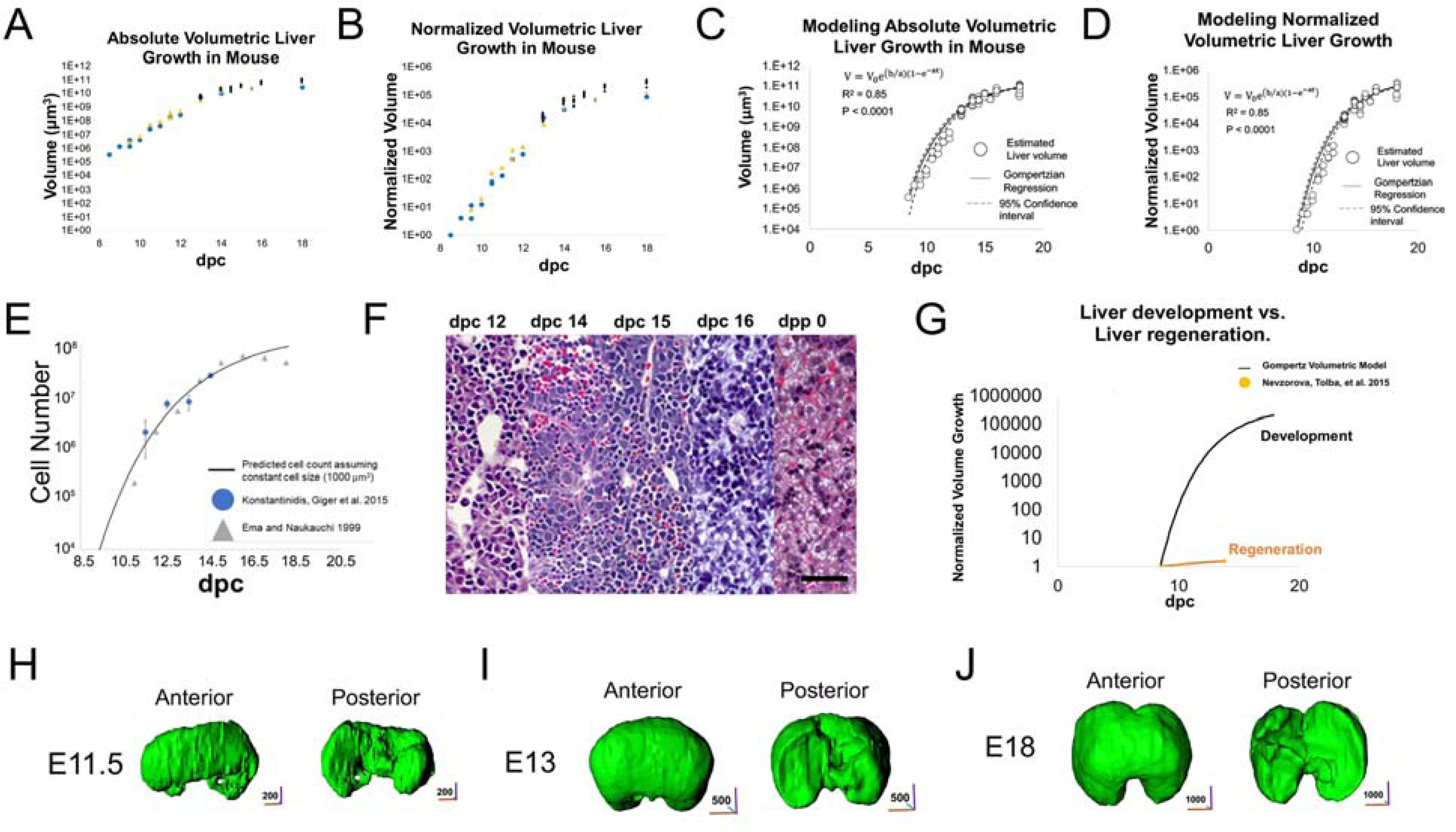
A quantitative model of volumetric growth of the murine developing liver. Since images demonstrated rapid liver bud growth, volumetric growth was quantified from E8.5-11.5 (embryonic stage) and E12-18 (fetal liver stage). Volumetric data was obtained from 3D reconstruction of tissue slices, MRI, optical projection imaging and liver weights. E8.5-E18 (n=1, E8.5; n=1, E9; n=3, E9.5; n=2, E10; n=3, E10.5, n=2, E11, n=3, E11.5; n=2, E12; n=32, E13; n=13, E14; n=11, E14.5; n=10, E15; n=35, E15.5; n=10, E16; n=12, E18). An observed 10^5^-fold increase in liver volume from E8.5 to E18. A) Absolute volume of the reconstructed 3D liver murine liver. Plotted points are identified in different colors based on source of data: optical projection (yellow), magnetic resonance imaging (gray), tissue section (blue), micro-CT (green) and liver weight (black). B) Relative liver volume was calculated by normalizing to the liver embryo volume at E8.5. Plotted points are identified in different colors based on source of data: optical projection (yellow), magnetic resonance imaging (gray), tissue section (blue), micro-CT (green) and liver weight (black). C) Absolute liver volume changes are modeled using a non-linear regression Gompertzian model of tissue growth and fitted to the equation 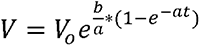. Assuming 95% confidence interval), *V_o_*=1.0E-72; a = 0.3069, 0.3245; an d b = 58.16, 61.40; RMSE = 11.42 x 10^9^. R^2^ =0.85, and P < 0.001. D) Relative liver volume was calculated by normalizing to the corresponding embryo volume. Changes are modelled using a non-linear regression Gompertzian model of tissue growth and fitted to the equation 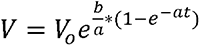. Assuming 95% confidence interval, *V_o_*=2.9E-78, a = 0.3069, 0.3245; and b = 58.16, 61.40; RMSE =3.29 x 10^4^. R^2^ = 0.85 and P < 0.001. E) Cell count estimated with the previously derived volumetric Gompertz model and assuming a fixed cell volume of 1000 µm^3^. Cell count data included from Konstantinidis et al. as blue circles. Each marker represents the mean and standard deviation of at least 5 fetal livers for each time point. Additional data also included from Ema and Naukauchi as gray triangles. Each marker is the mean of 7 or 8 fetal livers except dpc 11, which is the average of 31 fetal livers. F) H&E Stains of a mouse fetal liver at dpc 12, 14, 15, 16, and dpc 0(Ayres-Silva Jde, Manso et al. 2011). Scale Bar = 30 µm. G) A comparison of the relative volumetric growth for the Gompertz model and liver regeneration data from a partial hepatectomy (Nevzorova, Tolba et al.). Data was normalized to E8.5 in the Gompertz model and lowest liver weight after partial hepatectomy. H) 3D reconstruction of the liver bud at E11.5, E13, and E18 (green). Anterior and posterior views of the developing liver.

**Table IV:**
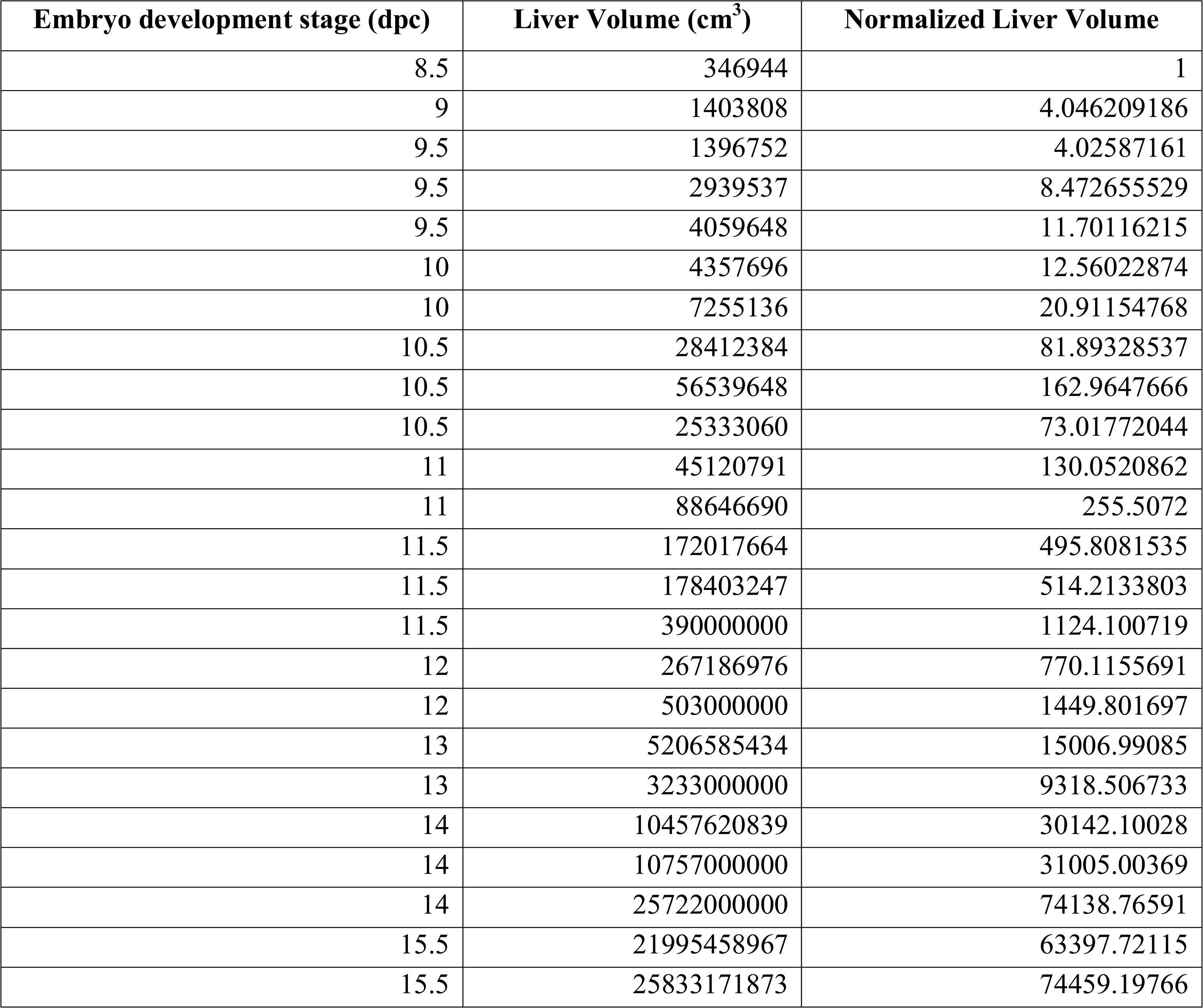
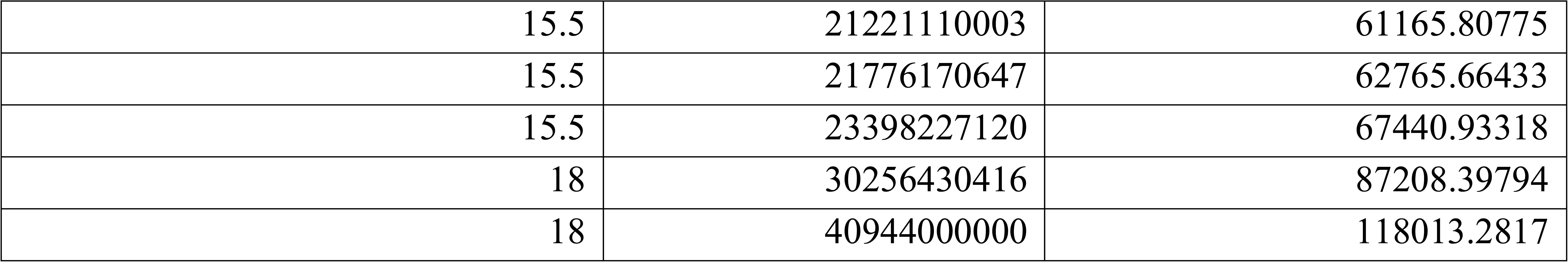
Liver Volume Estimation.

### 3D imaging of STM-epithelial interactions and growth during murine liver bud development

There is a renewed interest in mesenchyme interactions during liver organogenesis (Lotto, Drissler et al. 2020). Since mouse genetic studies confirm {Han, 2020 #125};{Asahina, 2011 #126}STM plays a large role in liver growth, we visualized interactions, with a variety of views, between STM and liver at E8.5 (**Figure 5A-C, Supplemental Video 6)**, E9.0 (**Figure 5D-F, Supplemental Video 7),** E9.5 (**Figures 5G-I, Supplemental Videos 8**), and E 10.0 (**Figures 5J-K, Supplemental Video 9**). Throughout E8.5-10, we found that in all cases STM envelops the liver bud, particularly at E9-10. We also observed that STM is not present on the dorsal part of the liver bud (**Figures 5I and 5L**). Finally, the STM appears to increase in size between E8.5-E10. We quantified the relative and absolute growth of the STM. We observed an 8.8 x 10^2^-fold increase in STM volume, as compared to 8.06 x 10^2^-fold increase in liver bud volume, over the same time period (**Figure 5A-B**). Overall, the data demonstrated that the STM mirrors the growth pattern of the mouse liver bud between E8.5-E10, and continues to envelop the liver bud with time.

**Figure 5.**
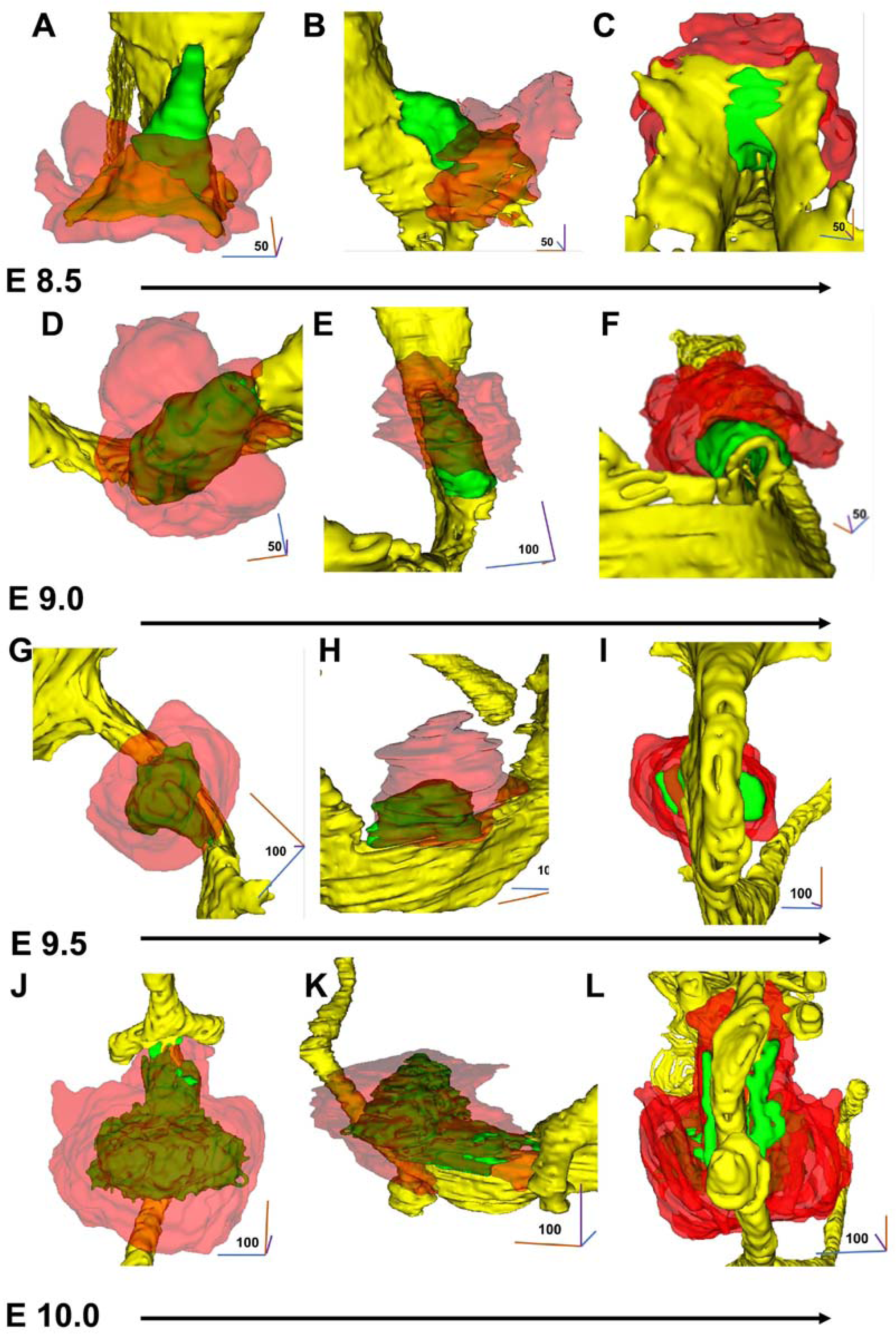
Visualization of 3D reconstructed murine liver bud and septum transversum mesenchyme (STM) interactions. The STM has a key role in liver development. To visualize liver bud-STM interactions during early liver bud morphogenesis, the liver bud, the gut tube, and the STM were visualized together during early liver bud development. This was performed as described in Figure 1. Each time point based on one specimen. A) 3D reconstruction of the liver murine liver bud and STM, at E8.5. Ventral view. Liver bud (green), STM (red). Visualization of the STM at E8.5 demonstrates that it covers the inferior ∼ ½ of the E8.5 liver bud. B) Same as A) except right lateral view. The STM extends ventrally for a distance similar to the dorsal-ventral thickness of the liver bud. C) Same as A) except inferior view. The width of the STM in the lateral direction is about ¾ of the lateral thickness of the liver bud. D) 3D reconstruction of the liver murine liver bud and STM, at E9.0 Superior, ventral, lateral view. the STM extends laterally for a length equal to ¾ times the lateral (smaller) diameter, and covers approximately 80% of the liver bud. E) Same as D) except ventral, right lateral view, demonstrating the STM extends laterally for a length equal to ¾ times the lateral (smaller) diameter, and covers approximately 80% of the liver bud. F) Same as E) except ventral, inferior (anterior) view. This demonstrates how the STM contacts the liver bud directly. G) 3D reconstruction of the liver murine liver bud and septum transversum mesenchyme, at E9.5 Ventral, lateral view. The STM extends laterally about ½ times the diameter of the liver bud, and fully surrounds the liver bud. This suggests that the STM grows, remodels with the liver bud, and potentially primes the liver for growth. H) Same as G) except right lateral view. The STM appears to be as thick as the liver bud when viewed laterally. I) Same as G) except dorsal view. The STM extends laterally, for a distance of 1/3 times the lateral width of the liver bud. J) 3D reconstruction of the liver murine liver bud and STM, at E10.0 Ventral, lateral view. Compared to E9.5, the lateral STM thickness has increased. K) Same as J) except left lateral view. The STM has diminished in the ventral direction. L) Same as J) except posterior view. The STM has diminished in the lateral direction.

### 3D imaging of migrating cords and sheet-like growth in the developing human liver bud

To our knowledge, there are few if any studies of human liver bud growth. Imaging of the human 3D reconstructed liver demonstrates extensive lateral growth on days 28 and 33 (**Supplementary Figure 7A-C**), and by day 33, we found that the liver epithelium (green) was indistinguishable from the STM (**Supplementary Figure 7C**). The day 25 liver bud demonstrates numerous 3D cellular projections consistent with hepatic cords in multiple directions (**Figures 6A-C, Supplemental Videos 10-11**). Interestingly, we again notice here not only narrow hepatic cords but also prominent sheet-like structures at multiple levels, particularly visible in anterior-lateral-inferior and anterior-lateral-superior views (**Figures 6A-C**). These cords and sheet-like structures were not due to artifacts of 3D reconstruction. For example, they are not present in the gut tube images of the same reconstructed images, and data was once again stacked, but not interpolated between tissue sections. These cords and sheets continue to be present at Day 28 (**Figures 6D-F, Supplemental Videos 12-13**) in multiple views, with a large sheet present in the ventral direction (**Figure 6E**) with clear layering or stacking of sheet-like projections (**Figure 6F**). By Day 33, the liver surface has smoothened and greatly enlarged (**Figures 6G-H, Supplementary Figures 7-8, Supplemental Videos 14-15**). At this time, the liver demonstrates a 3D ellipsoid shape with a lateral axis of ∼3500 µm and ∼1000 µm in the cranial/anterior direction, indicating extensive lateral growth. We imaged inside the 3D reconstructed liver to analyze interconnections, and observed portions of the liver in enclosed spaces, which appear as green as enclosed spaces (**Figure 6I, Supplementary Videos 16**). We quantified liver growth between day 25 and day 294 using the volume estimations from these 3D reconstructions, as well as published liver volume data (Szpinda, Paruszewska-Achtel et al. 2015) and liver weight data (Man, Hutchinson et al. 2016). We performed a non-linear regression analysis with this data to fit a Gompertz curve (R^2^ = 0.84, P < 0.001, RMSE = 3.08 x 10^6^) (**Figure 6J**) and for normalized to day 25 (**Figure 6K**) (R^2^ = 0.84, P < 0.001, RMSE = 1.04 x 10^3^). The data demonstrates a 2.41 x 10^3^–fold change in volume between day 25 and day 50, which is less than the 1.1 x 10^4^–fold change we measure from E9-E14 during mouse development (**Compare Figure 6K with Figure 4B**).

**Figure 6.**
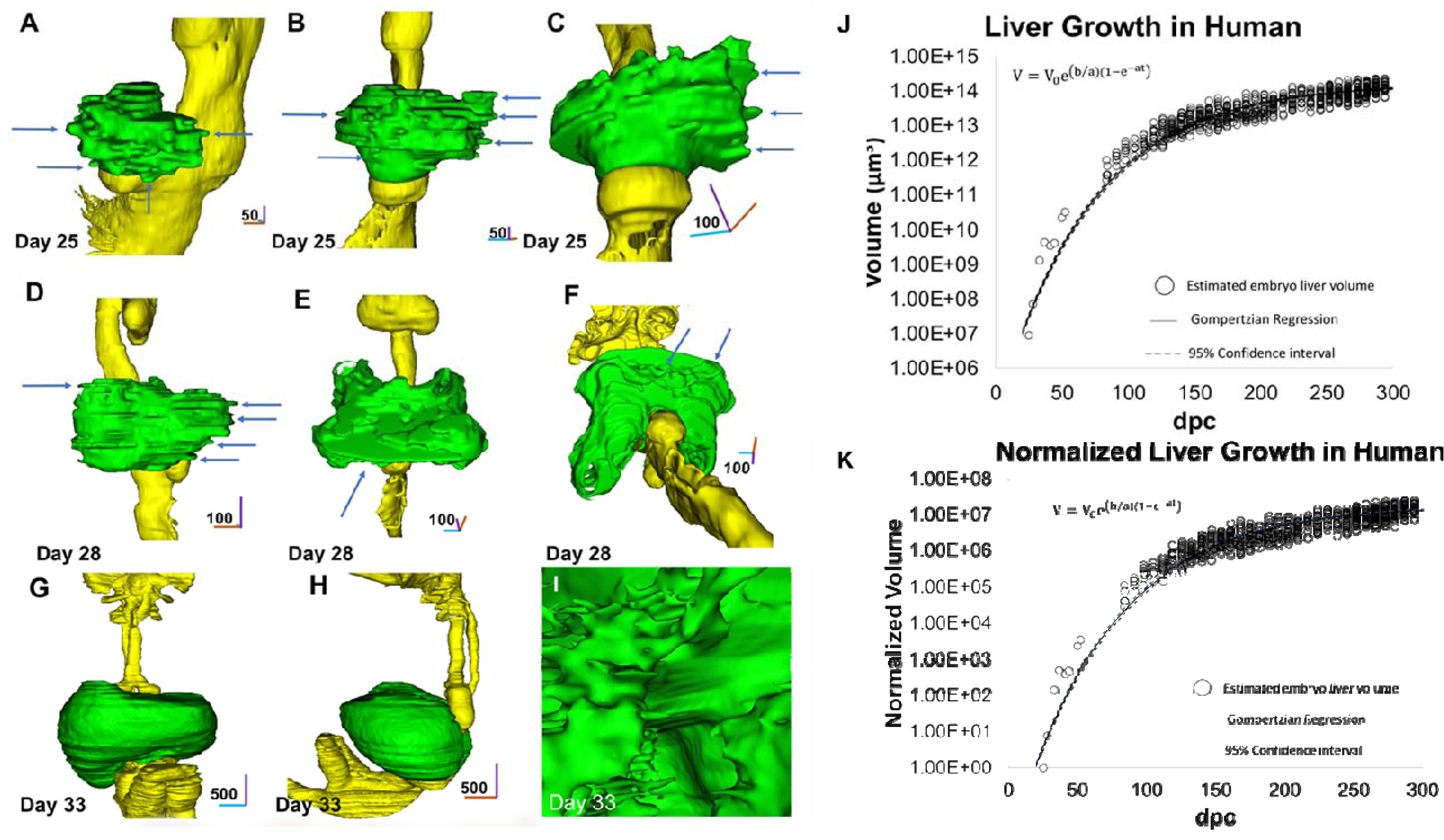
3D reconstruction and visualization of the developing human liver bud. To understand dynamics of liver bud morphogenesis, the human liver bud was imaged in 3D similar to the procedure used to visualize the mouse liver bud. 3D datasets, consisting of cleared hematoxylin and eosin (H&E) tissue sections from the 3D Atlas of Human Embryology were used. Datasets were reformatted, imported, segmented, and reconstructed in 3D. Ventral views are shown. Each time point based on one specimen. Gut tube (yellow), the liver bud (green). Sample per timepoint are n=9, Days 25-56, n = 930, Day 125 - 294 A) 3D reconstruction of the human liver bud on day 25. Arrows demonstrate hepatic cords (arrows), and sheet-like growth appears B) Same as A) except ventral view. Lateral hepatic cords (arrows) can be visualized, and sheet-like growth is again observed. C) Same as A) except, inferior, left lateral, ventral view. Lateral hepatic cords can be visualized (arrows), and inferior view enables observation of sheet like growth. In some cases, sheets of cells are bounded by hepatic cords. Layers of sheets can be clearly visualized. D) 3D reconstruction focused of the human liver bud on day 28. Lateral view. Arrows depict migrating cords. Although larger than day 25, the liver bud demonstrates sheet-like growth and hepatic cords. E) Same as D) except anterior, dorsal view. Arrow depicts tissue which wraps around anatomical structures. Dorsal side does not demonstrate appreciable hepatic cords. F) Same as D) except ventral view. Arrows depicting sheet of migrating cells extending medially and laterally. G) 3D reconstruction focused of the human liver bud on day 33. Ventral view. Image demonstrates that the liver has enlarged and is smoothened, with no hepatic cords or cell sheets are present. H) Same as G) except lateral view. Again, no hepatic cords or hepatic sheets are present. Inside view of the 3D reconstruction focused of the human liver bud on day 33. I) The green images demonstrate numerous enclosed green structures and trabeculation, all of which are in fact empty spaces within the 3D constructed liver. Interconnections can be seen inside the liver as the green portions are enclosed voids. J) Absolute human liver volume changes are modelled using a non-linear regression Gompertzian model of tissue growth and fitted to the equation 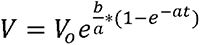. Assuming 95% confidence interval, whereby *V_o_* = 50,000, a = 0.014, 0.015; b = 0.31, 0.33; P < 0.001, R^2^ = 0.84 and RMSE = 3.06 x 10^6^ K) Relative human liver volume was calculated by normalizing to the embryo liver volume at day 25. Changes are modeled using a non-linear regression Gompertzian model of tissue growth and fitted to the equation 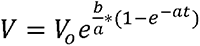. Assuming 95% confidence interval, whereby *V_o_*=0.0056, a = 0.014, 0.015; b = 0.31, 0.33; P < 0.001, R^2^ = 0.84 and RMSE = 1.04 x 10^3^

### 3D imaging of STM-epithelial interactions during human liver bud development

We reconstructed the liver bud together with STM to image their interactions. We analyzed several views on Day 25 (**Figures 7A-E, Supplemental Videos 17-18**) and Day 28 (**Figure 7F-H, Supplemental Videos 19-20)**. The human liver bud at Day 25 demonstrates that STM envelops the liver bud and extends all major directions (**Figures 7A-E, Supplemental Videos 17-18**). The human liver bud at Day 28 demonstrates that STM has grown compared to Day 25, however, visually is has now changed, since it is now absent on several surfaces of the liver. The reason is it absent is because cells of the STM have been absorbed into the liver epithelium and cannot be distinguished (**Figures 7F-H).** We quantified the STM growth compared to the liver growth on Day 25 and Day 28. The STM size increased by 0.54-fold, while the liver increased by 6.8-fold **(Figure 7I**).

**Figure 7.**
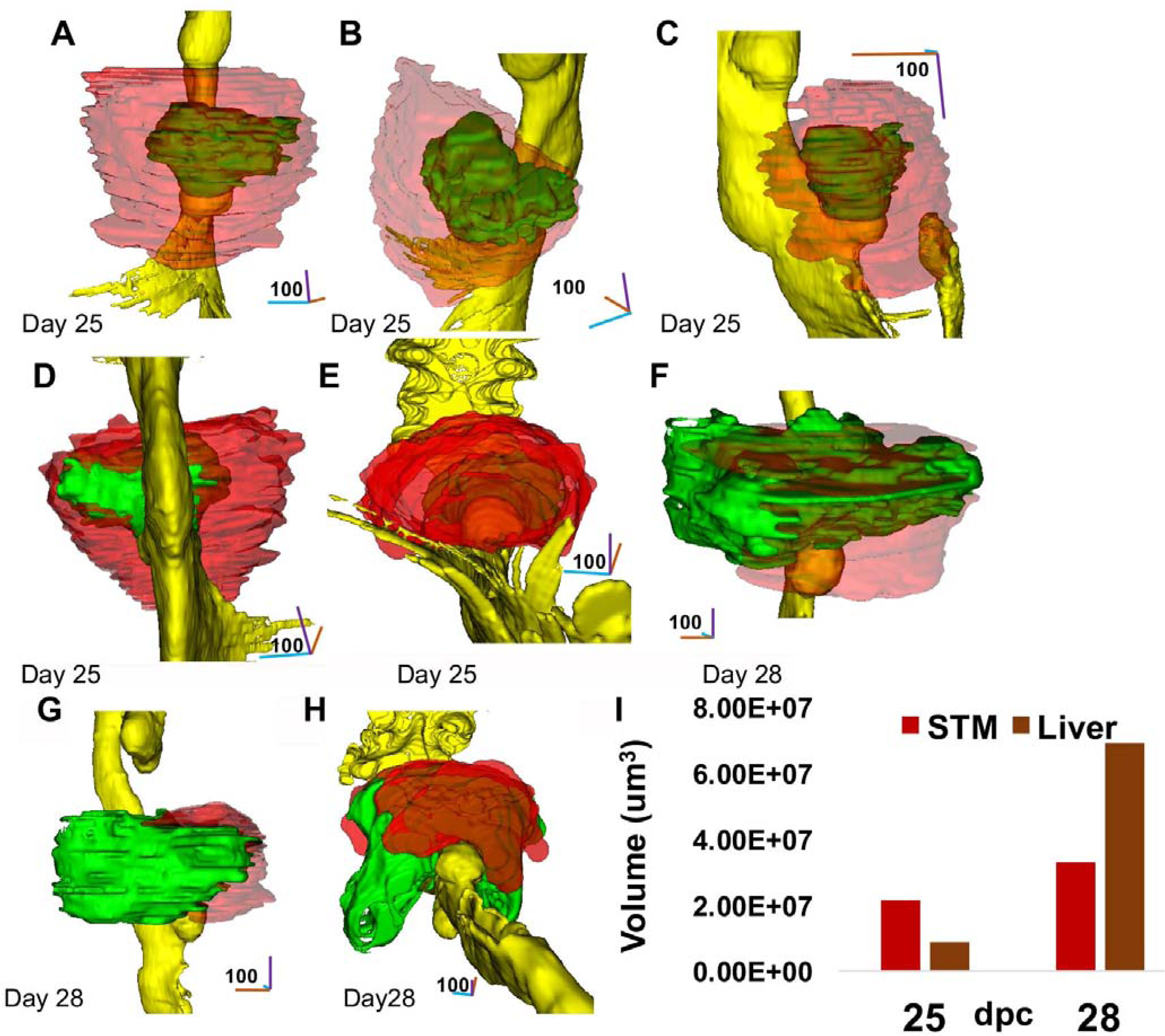
Visualization of 3D reconstructed human liver bud and septum transversum mesenchyme (STM) interactions. To understand the role of the STM in human liver bud morphogenesis, human liver bud was again imaged in 3D as in Figure 6, except the STM was added to the images. 3D datasets, consisting of cleared hematoxylin and eosin (H&E) tissue sections from the 3D Atlas of Human Embryology were used. Datasets were reformatted, imported, segmented, and reconstructed in 3D. Ventral views are shown. Each time point based on one specimen. Ventral, lateral view. The STM model was rendered at 50% transparency. Each time point based on one specimen. A) 3D reconstruction of the human liver bud and STM, at Day 25. Ventral view. Liver bud (green), STM (red). At this stage, the STM completely envelops the liver. The human liver bud is surrounded by the STM, extending about ½ times the width and height of the liver bud. The posterior is not fully covered. B) Same as A, except superior, lateral view. The STM continues to envelop the liver in all directions. C) Same as A, except right, lateral view. The STM envelops the liver. D) Same as D, except dorsal view. Portions of the liver bud are free from STM, in the posterior portion of liver bud. E) Same as A, except inferior view. STM envelops the liver bud, and appears to form a tail-like structure inferiorly. F) 3D reconstruction of the human liver bud and STM, at Day 28. Ventral, lateral view. The STM is observable superiorly and inferiorly, but within the sheet of cells within the center of the liver. G) Same as F, except right, lateral view. STM is not observed at this stage. H) Same as F, except inferior view with the ventral surface facing up. STM is not observed on the dorsal side of liver but is present on the ventral side. I) Quantification of human STM and liver bud volume during human liver development on days 25 and 28. The liver growth, in terms of volume, has overtaken the STM by day 28, unlike the mouse.

### Epithelial-mesenchymal interactions in an human stem cell-derived organoid model of human liver bud

To functionally determine the role of mesenchyme in embryonic liver bud growth (E8.5-E11), we developed an *in vitro*, hPSC-derived hepatic organoid model which models collective migration from the E8.5 liver bud into the STM, based on our recent study modeling STM and collective migration from cell lines (Ogoke et al. 2020). This model employed hPSC differentiation towards hepatic progenitors, engineering of hPSC-haptic progenitors-derived organoids, and STM modeling with organoids submerged in MG bearing human fibroblasts (HFF) (**Figure 8A-B**). Gene expression analysis demonstrated significantly increased transcriptional levels of alpha-fetoprotein and albumin (**Figure 8D**), suggesting day 12 hPSC-hepatic progenitors were specified along the liver lineage. Organoid culture in mesenchyme bearing MG + HFF demonstrated distinct cellular protrusions consistent with 3D collective migration that was not present in the absence of HFF (**Figure 8E**). Image analysis demonstrated significant changes in circularity (morphogenesis) and protrusions (3D Migration) (**Figure 8F**). Overall, this functional experimental model supports the liver bud imaging data in terms of the role of the STM in boosting morphogenesis and migration during embryonic liver development.

**Figure 8.**
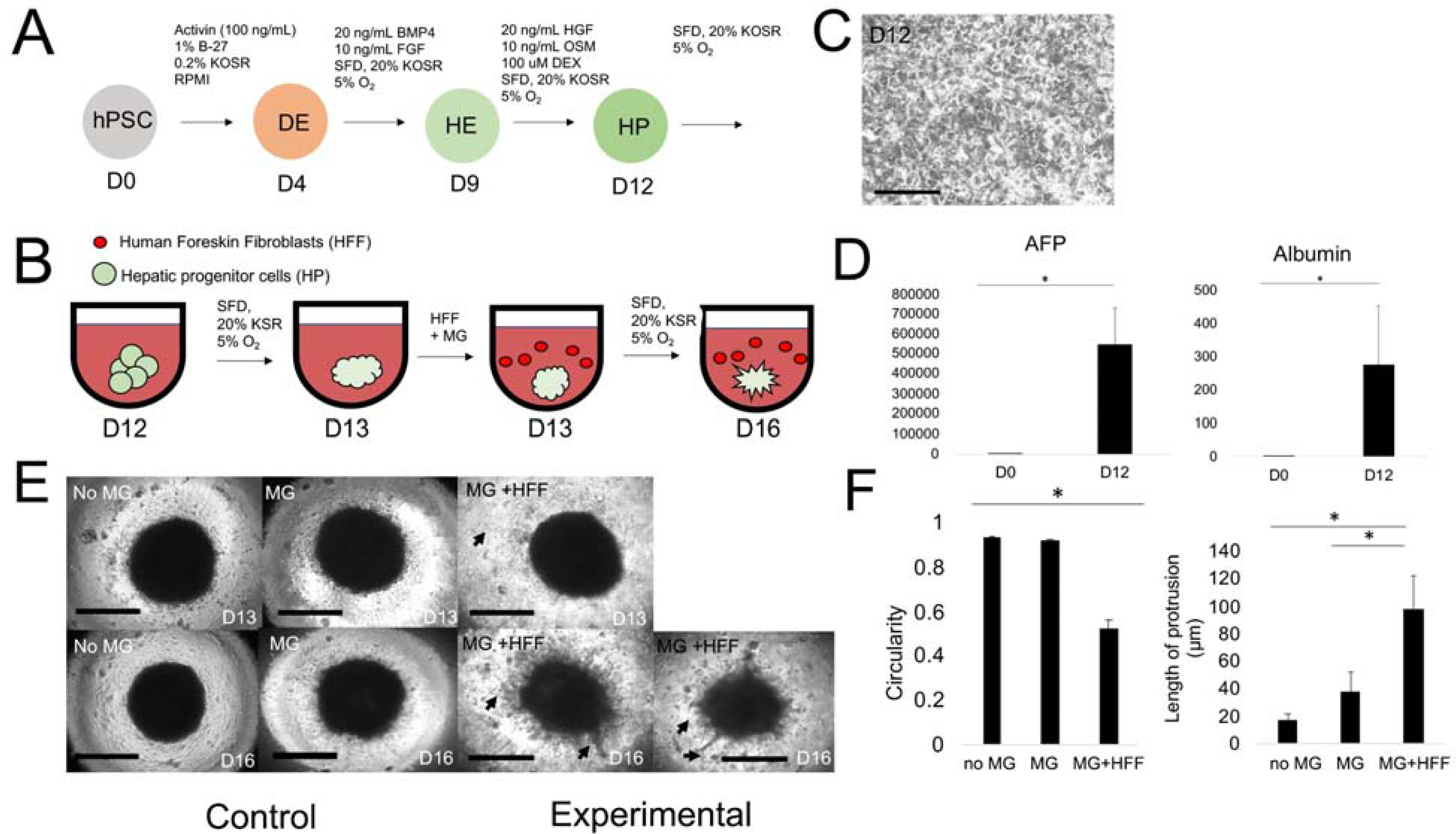
Epithelial-mesenchymal interactions in an human stem cell-derived organoid model of human liver bud. A) hPSC derived hepatic progenitor cells are differentiated using our illustrated scheme in from hPSCs to definitive endoderm (DE). Subsequently cells are directed towards hepatic lineage using BPM4 (20 ng/mL) and FGF (10 ng/mL) to form hepatic endoderm (HE). Cells are then exposed to Hepatocyte growth factor (HGF) - 20 ng/ml, Dexamethasone (DEX) - 100 µM, and Oncostatin M (OSM) - 10 ng/ml, for brief hepatic lineage maturation for form hepatic progenitors (HP). Medium is a nutrient rich medium (SFD) with 20% Knockout serum(defined serum), and experiments at under hypoxic conditions. B) Schematic of 3D organoid formation using hPSC hepatic progenitor cells. C) Morphology of 2D hepatic progenitor cells after Day 12 induction, Scale bar = 200 µm. Compact organoids form in 24 hours and can be subsequently embedded in Matrigel containing fibroblasts. D) qRT-PCR analysis of Day 12 hepatic progenitor cells demonstrating significant increases in alpha-fetoprotein (AFP) and Albumin (ALB) expression. Plotted is relative fold change normalized to GAPDH, (AFP, day 0 vs day 12, n=3, P =0.0069); (Albumin, Day 0 vs Day 12, n=3, P = 0.054). E) Images of hepatic organoids modeling liver bud in suspension culture, matrigel embedded culture, and Matrigel (MG) + human foreskin fibroblasts (HFF) embedded culture. There was significant migration in the MG+ HFF culture condition as compared to the no MG and MG no HFF from D13-D16. Scale bar = 300 µm. F) Analysis of liver bud organoid 3D culture, there is significantly less circularity of the central organoid of liver bud model in the Matrigel+fibroblast condition (n=3, P=0.0033). Subsequent characterization of the length of the protrusions from the central organoid demonstrated significant increases in the Matrigel+fibroblast culture condition over controls (MG+HFF vs. 3D sus, n=3, P = 0.0081); (MG+HFF vs. MG, n=3, P = 0.009). Plotted is mean ± SD. Significance defined as P ≤ 0.05.

## DISCUSSION

Scientists are particularly interested in modeling liver organogenesis *in vivo* for basic science, disease modeling, drug development and discovery, and therapeutic purposes, but many questions remain regarding murine and human liver organogenesis. An important aspect of liver organogenesis is, how does the liver become the largest internal organ in only 10 days of mouse fetal development? Both the underlying morphogenetic mechanisms involved (Koike, Iwasawa et al. 2019) and the role for intercellular, epithelial-mesenchymal signaling in cell migration (Lotto, Drissler et al. 2020) are not well understood. To answer these questions, we focused on serial, high resolution, 3D imaging and quantitative modeling of liver bud morphogenesis in mouse and humans, as well as functional experiments with hPSC-derived hepatic organoids. We imaged, at high resolution, 3D hepatic cord formation, mesenchymal-liver epithelial interactions, and embryonic and fetal liver growth in both mouse and humans. This enabled imaging of both murine and human hepatic cords in 3D, for the first time, and comparison of structural differences. We applied a mathematical modeling approach to develop a predictive model of growth, and used this to model cell kinetics and compare liver bud growth versus liver regeneration. Further, our model demonstrated previously underappreciated exponential liver growth in early embryonic liver growth and our growth curve matched embryonic to fetal liver growth and was validated by existing liver imaging and fetal liver weight data. We identified morphogenetic features associated with growth, including 3D hepatic cord formation, sheet-like growth, and lateral growth. We also demonstrate that the STM increases in size during early fetal liver growth, is highly interconnected with liver epithelium, and that it may have local effects on growth and remodeling, and our functional experiments with hPSC-hepatic organoids and STM modeling support these findings. Overall, we provide several novel findings that contribute to an improved understanding of liver organogenesis.

Embryonic and fetal liver growth establishes the early tissue architecture of the hepatic, biliary, and vascular components of the liver. Our data suggests that the mouse liver undergoes an overall 8.74 x 10^4^-fold-increase in volume **(Figure 4B**) between E8.5 to E18. Further, we found a 7.5 x 10^3^ change in volume from E9-E14, which corresponds to a 2.5 x 10^4^–fold change in volume between day 25 and day 56 in humans (**Figure 6K**). This is in contrast to our data at later stages of liver growth which demonstrates an approximate 7.3-fold increase in volume (**Figure 6K**) (Szpinda, Paruszewska-Achtel et al. 2015) between day 126 and 210 (18 weeks to 30 weeks). We also utilized this model to estimate cell number over time (**Figure 4E**), and demonstrate the exponential growth of liver bud compared to the more linear growth of liver regeneration (**Figure 4G**). These studies highlight the uniqueness of early liver bud growth. Our data compares favorably to studies of fetal liver proliferative capacity that demonstrate embryonic day 14 (E14) fetal rat hepatocytes (rat F-Heps) are still proliferating after 6 months (or the equivalent of 12 human years) with full rat liver repopulation (Dabeva, Petkov et al. 2000). Further, studies of rat E18 F-Heps or phenotypically isolated rat E19 fetal hepatic progenitors were shown more recently to repopulate ∼1/3 the liver mass in acute and chronic models (Boylan, Francois-Vaughan et al. 2017);(Bin, Ma et al. 2012). Although the mechanisms by which the exponential increase in liver bud growth occurs are not well understood, our study suggests that that fetal hematopoiesis or cellular hypertrophy are not fully responsible. For example, the most rapid growth occurs initially, from E8.5 to E10.0, a time when fetal liver hematopoiesis is not believed to have fully commenced (Lotto, Drissler et al. 2020), although at later stages from E10.0-E18, we still observed rapid growth which could be attributable to hematopoiesis. Further, our tissue section analysis from the literature demonstrates no clear evidence of cellular hypertrophy (**Figure 4F**). Overall, we feel our quantitative modeling approach can lead to further development of improved models of cell kinetics, which match existing liver cell kinetic growth models in the literature (**Figure 4E**).

Further studies to enhance our understanding of growth may involve careful measurements for the development of population kinetic models, including modeling of the proliferative capacity of multiple cell types, cell death pathways, and cellular metabolism. Improved models can also integrate potentially more complex mechanisms involving suppression of proliferation, cell death, and organ size control that regulate growth at fetal liver growth, and potentially be used to understand organoid growth.

A key finding is that we report high resolution imaging of the 3D hepatic cords in humans and mice. Interestingly, we observe hepatic cords at an earlier stage in mice (E9.0) than previously reported (E9.5). This is likely because our technique has a higher spatial resolution than tissue sectioning, and samples the entire volume of the liver, rather than a single section. Our 3D data agrees with existing data and confirms that these cords migrate to form the trabeculated liver by E10.0 (**Figure 3H**). It remains to be determined how the 3D hepatic cords form the trabeculated liver, as this occurs very rapidly between E9.5 and E10. Thus, improved high temporal and spatial resolution imaging is required between these time points to further elucidate these mechanisms. Our data suggests that the human liver bud generally has more hepatic cords than the mouse, and more predominantly in the lateral directions. This could be because the liver grows even larger in humans, and lateral growth is a key mechanism. Further, we observed these hepatic cords merging into sheets in the human liver, prior to trabeculation, in both mouse and humans. We speculate that this additional “sheet morphogenesis” step may be critical for exponential growth which occurs in the mouse and human liver. Previous studies have used the term “sheet-like” growth in early fetal liver cultures (Hussain, Sneddon et al. 2004), and the “hepatocyte sheets” have been used to describe mature liver architecture. While hepatic cords have been described to form hepatic sheets that flank the sinusoids, it is unclear if “sheet-like morphogenesis” has been identified or imaged. From our images, it is also unclear if there are layers of mesenchyme between the observed sheets, which would be an interesting subject of a future study.

Our analysis demonstrates the importance of interactions between the emerging liver epithelium and the STM. Interestingly, our murine data indicates the STM grows at a nearly identical rate to the liver bud between E8.5-E10, and the absolute volume of the STM slightly larger than that of the liver bud from E8-E10. These two facts are not obvious from the approach of traditional tissue sections. While STM expresses inductive signals like BMP, the expanding STM volume indicates the potential complex role of the STM and reciprocal signaling between the hepatic endoderm/hepatoblasts. This is consistent with the role of GATA6 and GATA4 within the STM, in which GATA6 and GATA4 knockout via tetraploid blastocyst complementation demonstrates loss of STM mass and inhibits early liver bud development (Watt, Zhao et al. 2007);(Zhao, Watt et al. 2005). Our analysis indicates that large portions of the STM are in a remote location from the liver bud, raising questions of what their role is, because if they were secreting substances to liver epithelium, this would take place over very long distances. Further, human liver bud formation suggests that after sheet-like growth, the STM has been obliterated and internalized laterally, while not in the anterior directions. This strongly suggests the STM may play a role in sheet formation and incorporation of the STM into the liver (**Figure 7H**). However, we can only speculate regarding the mechanisms by which the STM may accomplish this. Another potential role of the STM is in contributing to liver bud remodeling, and not just growth. Ultimately, our objective was to provide a critical analysis of liver bud development in murine and human models. It’s quite possible that STM interactions are not equally important in human and mouse development. Here one of our novel findings is that the STM grows with the liver, which may be obvious, but was not previously stated in liver bud literature. Therefore, our modeling points to reciprocal interactions between STM-liver bud interactions that when recreating the process.

Our visualization and modeling approach can enhance existing approaches to model 3D liver organogenesis *in vitro*. Based on our data, it appears that current *in vitro* models may not fully accurate 3D liver bud morphogenesis. With regards morphogenesis we identified that rapid alterations in liver shape, orientation and size (mass) take place, and our data is consistent with recent data indicating that large population of migrating cells at E9.5 (Lotto, Drissler et al. 2020). While we recognize that cues regulating these phenomena might be cell intrinsic, the adjacent tissue, the STM also rapidly changes in parallel with the liver. Thus, our data provides strong evidence for a supportive role of mesenchyme in facilitating liver bud migration and morphogenesis, which might be valuable for *in vitro* modeling. To support this idea, our liver bud *in vitro* models, based upon recent work from our lab (Ogoke et al. 2020), strongly support the notion that mesenchymal cells support early migration from organoids, which with further research, could be used to potentially expand liver tissue. Here we have recreated these models with hPSC-derived hepatic organoids **(Figure 8E-F)** which demonstrate clear evidence of 3D migration. Thus our imaging data and our *in vitro* models suggest that migration is a critical step in the exponential growth we observed during liver growth, which improve current generations of hepatic organoid models, which we have recently reviewed (Ogoke, Maloy et al. 2021). While our data strongly suggests mesenchyme bearing HFF is required for driving migration/tissue morphogenesis, additional functional assays, mesenchyme cell perturbations, and dynamic transcriptomic and proteomic analysis of 3D tissue models will aid in further elucidating mechanisms in recapitulating embryonic liver growth.

For our cell kinetics analysis, we estimated cell size to calculate an approximate cell number over time. To make this estimation, we estimated the volume of all cells (hepatic and endothelial progenitors, stellate cells, hematopoietic stem cells, immune cells) to be 1000 µm^3^, i.e., average volume of human cells. In terms of the cell volume estimation, it is also important to note that there exists potential non-cell-bearing, intra-organ space in the growing liver. In fact, putative liver stem cells have a diameter of ∼10 µm so they can be estimated to accommodate ∼1000 µm^3^ of space (Darwiche and Petersen 2010). Further complicating the cell volume calculation, studies demonstrate that the intercellular distance between hepatoblasts is believed to be a 4-5 µm. Another important point is that mature adult hepatocytes have larger volumes (∼3000 µm^3^), although the cell volumes estimated here refer to the embryonic and fetal liver (Chen, Soto-Gutierrez et al. 2018). Since hepatoblasts are smaller than hepatocytes, the average size of the hepatic cells likely increases with time. Further, endothelial cells are significantly smaller than hepatoblasts (Pandey, Nour et al. 2020); (Marguerat and Bahler 2012)and may have higher packing densities and less intercellular space than hepatocytes. Therefore, although the average cell volume can be more accurately calculated at each time point, we feel justified in estimating 1000 µm^3^ cell volume, accounting for all the complicating factors mentioned above. To calculate the exact cell volumes experimentally, one would have to collect embryonic and fetal livers daily between E8.5-E18, perform cellular digestion, perform staining and cell sorting, estimate cell size and cell fraction from flow cytometry data (or some other technique), and then calculate cell number over time. To our knowledge, the cell size throughout embryological to fetal liver development has not been well established. We have illustrated that there are minimal cell diameter changes during liver development from the literature **(Fig. 4F**). While it is experimentally feasible to calculate cell volumes over time, we felt this approach was outside of the scope of our study, and we have not determined the exact numbers here.

It is important to note that the finger-like projections and/or sheet-like structures we observe in both liver bud imaging and mesenchyme imaging studies could be confounded with artifacts of tissue sectioning or of 3D reconstruction. Here we discuss several problems pertaining to this issue and try to provide evidence that these factors are not responsible for generating the images that we have obtained. First, both mouse and human data sections were whole embryo sections (not at the organ level) and reconstructed with appropriate voxel size without the use of interpolation. In other words, the 3D slices were stacked on top of each other and at the correct sizes, but no data was created or interpolated between slices. Any smoothing functions in the software were not at the level of pixel size (µm) and therefore did not remove the sheet-like structures and protrusions that we observe. Error due to rotation of the slices is minimal, because the object was fixed and sectioned, and images were co-registered in the dataset itself. During sectioning of the tissue, there could be some physical displacement of the tissue, that when reconstructed would no longer remain, and this may be responsible for some of the layering effects that can be seen (**Figure 3C and Figure 6B,D)**. We would expect these errors to be on the scale of the sectioning blade itself. However, the sheet-like structures we observe in the human liver bud, are on the order, of 10-20 µm, while thin sectioning blades are on the order of 250 µm. A simple wrinkling effect can be observed in the mouse and human gut tube (**Figure 2B and Figure 6)** but sheet-like growth and formal protrusions are not observed in the area of the liver bud, indicating that these processes are likely unique to the liver. It is possible that liver tissue cuts with properties that are different from the gut tube, but these effects would appear more randomly, rather than as ordered structures. While the tissues were cleared before sectioning, this does not explain the effects we observed. Overall, we are comfortable that the findings that we observed are likely to be real. A good illustration of the high spatial resolution of the technique is the imaging of the E10 mouse liver (**Figure 2B, Figure 3F-3H**). It is very clear that these images are not due to artifacts. Another limitation of our imaging approach is in regards to the fact that STM cells that have integrated into the liver bud between E8.5 and E10. The trabecular pattern we identify within the liver bud at the onset of E8.5, represents the beginning of the primitive sinusoids. Although there is not an established method to quantify STM cells within the liver bud, we estimate it to be 5%, approximately the same percentage as in the adult liver.

One of the major weaknesses of our study was that there was only n = 1 mouse and n = 1 human for each time point analyzed for the 3D reconstruction of embryonic stages of liver bud development. This can only be improved by expanding the data set, which was outside of the scope of this study. The mouse databases we used each have between 1-4 mice for each condition tested, but only 1 data set per time point is available for download. Although we analyze morphological images of a single mouse at each time point, there are many sources of variation even if more mouse numbers had been used. There are many potential factors that cause variation in morphology as reported in the literature (Figure 3L-M), and it is unclear what is due to mouse variation versus technical variation (after tissue collection/dissection). A potential source of mouse variation may be the variation associated with somite-staging of embryological age (for example, E9.5 mice can have a variation in somite number). Further sources of variation may be due to differences in mouse littermates, differences between strains of mice, and phenotypic variation between individual mice. Another source of variability is technical variability. This technical variability could result from differences in mouse embryo isolation, dissection, angles of liver bud sectioning, tissue storage, tissue processing, sectioning, and staining. Because of this extensive variability between E8.5-E10, we feel that our data is in line with the rest of the literature. We feel that although performing more dissections is appropriate, a high amount of variation would be present and needs to be further addressed.

Our data from digital datasets (EMAP and 3D Embryo Atlas websites) was validated in our study using published imaging and liver weight data. In fact, we obtained tens to hundreds of other data points at later times of mouse development to validate. We employed mouse imaging data from CT, MRI, optical projection tomography, and ultrasound. It is important to note that MRI data at E9.5 was likely overestimated because we were unable to distinguish between liver bud epithelium and STM. In fact, 6 of the 9 MRI-measured liver volumes were higher than the 3D reconstructed volumes from tissue sections, but the trends that we measured were similar. The only other technique to get more reliable and quantitative measurement is by noninvasive imaging of the intact embryo *ex vivo* or *in vivo*. However, any removal of the liver bud would essentially destroy it. The problems with *ex vivo* imaging, assuming the embryo is no longer alive, is that embryo tissues will have lost tension, as well as blood flow, and organs will lose shape. Further, depending on the imaging technique, there can be many confounding factors to getting the high resolution images we have obtained and analyzed, and very likely the liver bud would lose any sources of potential contrast. The problem with *in vivo* imaging as the only option is that there will be motion artifact, due to both the mother and the embryo, which will cause a loss of spatial resolution, and more confounding layers like extraembryonic tissues (yolk sac) and structure(umbilical vein), that will cause most imaging techniques to lose considerable spatial resolution. Therefore, there cannot be a more quantitative and reliable technique, at a higher resolution, provided our tracing of the organ was accurate, using any existing imaging technology or technique. It is important to note that our approach was validated by our Gompertzian curve fits, which predict rapidly growing tissue initially, which also agrees with our manual measurements of data present in mouse atlas. A final limitation of the study is the relative assignment of pixels towards one versus another tissue group. We carefully reviewed the literature to make evidence-based predictions about the spatiotemporal location of the respective tissues, segmented by color. The gut tube was highlighted in yellow, the liver diverticulum as an outpocketing of the gut tube was highlighted in green. The STM of the liver bud was then highlighted in red. We acknowledge that there could be minor errors using this approach and that more exact methods could be used to confirm these results. However, these errors would not explain the regular structures that we observe and would only contribute minor errors to our volume calculations without changing the trends observed.

In summary, we feel we have contributed new information regarding liver growth, liver volumes, liver morphogenesis and liver-STM interactions at the earliest time points of liver development. However, further work and new tools are needed to elucidate molecular mechanisms by which hepatoblasts and STM interact to result in exponential liver growth. In addition, we will look to perform perturbation studies that explore cell-cell interactions as well as cell identity experiments that provide evidence for the types of cells present at each stage of liver development. With new clearing agents, whole embryos could be labeled with antibodies followed by wholemount imaging of the immunofluorescence. Multiple antibodies could be used to distinguish liver epithelium and the different cell types of the STM. This alternative approach would provide very high resolution and additional cellular information that has not yet been obtained.

## CONCLUSION

In conclusion, we have identified new aspects of liver growth, liver morphogenesis, and interactions between liver epithelium and STM. Newer techniques can be employed, like CT, MRI, photoacoustic imaging, and optical projection imaging, to improve datasets, and coupling 3D anatomical information to molecular information like gene expression, could be performed using techniques liked expansion microscopy (Chen, Tillberg et al. 2015). Another approach to understand these morphogenetic steps is whole embryo culture at the earliest stages. Overall, we feel understanding and imaging 3D these processes may lead to further insights about how the liver manages to greatly expand its mass while establishing an underlying complex architecture.

## ABBREVIATIONS

2D: two-dimensional
3D: three-dimensional
EMAP: emouse atlas project
STM: septum transversum mesenchyme
HEP: hepatocytes
FGF: fibroblast growth factor
BMP4: bone morphogenetic protein 4
FOXA2: forkhead box 2
Hex: homeobox protein
Prox1: prospero homeobox 1
Tbx3: T-box transcription box factor 3
EMT: epithelial to mesenchymal transition
TGFB: transforming growth factor beta
ARF6: ADP-ribosylation factor 6
VEGF-A: vascular endothelial growth factor
HGF: hepatocyte growth factor
IL-6: interleukin 6
HNF4a: hepatocyte nuclear factor 4 alpha
GATA6: GATA Binding Protein 6
GATA4: GATA Binding Protein 4
CT: computed tomography
MRI: magnetic resonance imaging

## SUPPLEMENTARY FIGURE LEGENDS

**Supplemental Figure 1. Segmentation process of murine liver development in whole embryo at E8.5.**

A) Coronal, black and white, hematoxylin and eosin (H+E) stain tissue section at E8.5 of mouse development, obtained from eMouseAtlas database, and corresponding segmented image. In segmented image, the gut tube (yellow), the liver bud (green) and STM (red) were segmented. A = Anterior, V = Ventral, P = Posterior, D= Dorsal. Arrows depict liver bud.

A) Same as A, except sagittal sections and corresponding segmented image.

**Supplemental Figure 2. Dynamic 3D imaging of murine liver development in whole embryo.**

A) Coronal, black and white, hematoxylin and eosin (H+E) stain tissue section at E9.0 of mouse development, obtained from eMouseAtlas database, and corresponding segmented image. In segmented image, the gut tube (yellow), the liver bud (green) and STM (red) were segmented. A = Anterior, V = Ventral, P = Posterior, D= Dorsal. Arrows depict liver bud.

B) Same as A, except sagittal sections and corresponding segmented image.

**Supplemental Figure 3. Segmentation process of murine liver development in whole embryo at E9.0.**

Same as Supplemental Figure 2 except E10.0. Scale bar is in microns.

**Supplemental Figure 4. Segmentation process of murine liver development in whole embryo at E10.0.**

3D reconstruction of the liver bud at E8.5, Transverse view. Scale bar is in microns.

**Supplemental Figure 5. Quantification of septum transversum mesenchyme (STM) volume during murine liver development.**

To measure the dynamics of STM growth, volumes were measured after loading segmented images of embryos. Relative volumes were normalized by taking the ratio to the corresponding embryo volumes. Each time point based on one specimen.

A) Ratio of the reconstructed 3D liver murine liver growth compared to the mouse embryo at E8.5-E10. STM growth mirrors the growth patterns of the liver bud.

B) Absolute volume of the reconstructed 3D liver murine liver growth at E8.5-E10. STM does not stay constant, but exhibits growth that patterns growth of the liver bud, including discontinuous growth.

**Supplemental Figure 6. 3D reconstruction and visualization of human liver development in whole embryo.**

A) Whole human embryo 3D reconstruction demonstrating reconstructed human embryo on Day 25. Each tissue of interest was identified and thresholded and traced on individual, transverse tissue slices using known cross sectional anatomy and pre-identified labels within the database. Gut tube (yellow), the liver bud (green) and STM (red). A = Anterior, V = Ventral, P = Posterior, D= Dorsal.

B) Same as A except Day 28 in human development.

C) Same as B except Day 33 in human development.

**Supplemental Figure 7. Ventral view of human liver development at Day 33.**

The liver bud (green) and gut tube (yellow) is viewed at the anterior position looking towards ventral. Scale in microns.

**Supplemental Figure 8. Dorsal view of human liver development at Day 33.**

Dorsal view of liver bud (green) and gut tube (yellow). Scale in microns.

**Supplemental Video 1. Murine liver bud at E8.5 in 360° rotation through sagittal plane.**

Ventral view of liver bud (green) rotating through sagittal plane. Units in axis are microns.

**Supplemental Video 2. Murine liver bud at E9.0 in 360° rotation through sagittal plane.**

Ventral view of liver bud (green) rotating through sagittal plane. Units in axis are microns.

**Supplemental Video 3. Murine liver bud at E9.5 in 360° rotation through sagittal plane.**

Ventral view of liver bud (green) rotating through sagittal plane. Units in axis are microns.

**Supplemental Video 4. Murine liver bud at E10.0 in 360° rotation through sagittal plane.**

Ventral view of liver bud (green) rotating through sagittal plane. Units in axis are microns.

**Supplemental Video 5. Interior view of murine liver bud at E10.0.**

Interior view of liver bud (green) through sagittal plane. Interior view is inverted as the structures (green) are voids within the liver bud and empty space is where the liver bud is solid. Units in axis are microns.

**Supplemental Video 6. Murine liver bud and STM at E8.5 in 360° rotation through sagittal plane.**

Ventral view of liver bud (green) and STM (red) rotating through sagittal plane. Units in axis are microns.

**Supplemental Video 7. Murine liver bud and STM at E9.0 in 360° rotation through sagittal plane.**

Ventral view of liver bud (green) and STM (red) rotating through sagittal plane. Units in axis are microns.

**Supplemental Video 8. Murine liver bud and STM at E9.5 in 360° rotation through sagittal plane.**

Ventral view of liver bud (green) and STM (red) rotating through sagittal plane. Units in axis are microns.

**Supplemental Video 9. Murine liver bud and STM at E10.0 in 360° rotation through sagittal plane.**

Ventral view of liver bud (green) and STM (red) rotating through sagittal plane. Units in axis are microns.

**Supplemental Video 10. Human liver bud at Day 25 in 360° rotation through sagittal plane.**

Ventral view of liver bud (green) rotating through sagittal plane. Units in axis are microns.

**Supplemental Video 11. Human liver bud at Day 25 in 360° rotation through sagittal plane.**

Ventral view of liver bud (green) rotating through transverse plane. Units in axis are microns.

**Supplemental Video 12. Human liver bud at Day 28 in 360° rotation through sagittal plane.**

Ventral view of liver bud (green) rotating through sagittal plane. Units in axis are microns.

**Supplemental Video 13. Human liver bud at Day 28 in 360° rotation through sagittal plane.**

Ventral view of liver bud (green) rotating through transverse plane. Units in axis are microns.

**Supplemental Video 14. Human liver bud at Day 33 in 360° rotation through sagittal plane.**

Ventral view of liver bud (green) rotating through sagittal plane. Units in axis are microns.

**Supplemental Video. 15. Human liver bud at Day 33 in 360° rotation through sagittal plane.**

Ventral view of liver bud (green) rotating through transverse plane. Units in axis are microns.

**Supplemental Video 16. Interior view of human liver bud at day 33.**

Interior view of liver bud (green) through sagittal plane. Interior view is inverted as the structures (green) are voids within the liver bud and empty space is where the liver bud is solid. Units in axis are microns.

**Supplemental Video 17. Human liver bud and STM at Day 25 in 360° rotation through sagittal plane.**

Ventral view of liver bud (green) and STM (red) rotating through sagittal plane. Units in axis are microns.

**Supplemental Video 18. Human liver bud and STM at Day 25 in 360° rotation through sagittal plane.**

Ventral view of liver bud (green) and STM (red) rotating through transverse plane. Units in axis are microns.

**Supplemental Video 19. Human liver bud and STM at Day 28 in 360° rotation through sagittal plane.**

Ventral view of liver bud (green) and STM (red) rotating through sagittal plane. Units in axis are microns.

**Supplemental Video 20. Human liver bud and STM at Day 28 in 360° rotation through sagittal plane.**

Ventral view of liver bud (green) and STM (red) rotating through transverse plane. Units in axis are microns.

## ETHICS APPROVAL AND CONSENT TO PARTICIPATE

Not applicable.

## CONSENT FOR PUBLICATION

Not applicable.

## AVAILABILITY OF DATA AND MATERIALS

All data generated or analyzed during this study are included in this published article and its supplementary files.

## AUTHOR CONTRIBUTIONS

OO: Obtained data, analyzed data, developed methodology for acquisition and analysis, wrote and approved manuscript.

DG: Obtained data, analyzed data, developed methodology for acquisition and analysis, wrote and approved manuscript.

TM: Obtained data, analyzed data, developed methodology for acquisition and analysis, wrote and approved manuscript.

CS: Obtained data, analyzed data, approved manuscript

SR: Obtained data, analyzed data, approved manuscript

SR: Obtained data, analyzed data, approved manuscript

NP: Conceptualized, acquired funding, investigated, supervised, wrote, edited manuscript, and approved manuscript.

## ACKNOWLEDGEMENTS

We thank Dr. Duncan Davidson and Professor Richard Baldock for their efforts in developing the eMouseAtlas Project which provided the datasets we used to perform our embryological analysis. We thank Professor R. Mark Henkelman, Dr. Jason P. Lerch, Dr. Brian J. Nieman, and Dr. John G. Sled for their work as part of the Mouse Imaging Centre and for providing us with additional image data sets for mouse development. We also like to thank Professor Bernadette S. de Bakker and the rest of the 3D Atlas team for providing us with human embryo data sets.

